# A novel mechanism of Euonymine inhibits in-stent restenosis through enhancing contractile phenotype of VSMCs by targeting AKT1 and p38MAPK

**DOI:** 10.1101/2021.10.29.466441

**Authors:** Li Zhang, Yi Ting Tao, Qin Hu, Ren Hua Yang, Jia Jia, Hao Nan Jin, Yong Zhao Yang, Yan Yang, Ming Yang Yu, Yu Ting Wang, Jia Ning Shi, Dan Yu, Guang Ping Tang, Jie Xu, Bao Xiong, Zhi Qiang Shen, Zhuo Yu, Hong Tao Qin, Peng Chen

## Abstract

This study aimed to examine the inhibitory effects of Euonymine on in-stent restenosis (ISR) after percutaneous coronary intervention (PCI) and oxidized low-density lipoprotein (ox-LDL)-induced proliferation, migration, and pro-apoptotic of vascular smooth muscle cells (VSMCs) in vitro, and its potential mechanisms. Euonymine is a monomer component extracted from Tripterygium hypoglaucum (Levl) Hutch. Using in vitro models of rabbit carotid balloon injury and porcine atherosclerotic coronary implantation, we confirmed that Euonymine inhibited ISR after PCI. Furthermore, Euonymine inhibited VSMC phenotypic transformation by targeting AKT1 to regulate the PTEN/AKT1/m TOR signaling pathway, with exertion of anti-proliferative, anti-migratory, and pro-apoptotic effects on ox-LDL-induced cell injury model. Additionally, the study demonstrated that Euonymine induced apoptosis of VSMCs via the p38MAPK-related mitochondria-dependent apoptotic pathway. Collectively, these findings indicated that Euonymine drug-eluting stents inhibited ISR after PCI by targeting AKT1 and p38MAPK to enhance the contractile phenotype of VSMCs to prevent intimal hyperplasia development. This provides insights into a potential therapeutic strategy involving the beneficial effect of Euonymine drug-eluting stent on ISR.

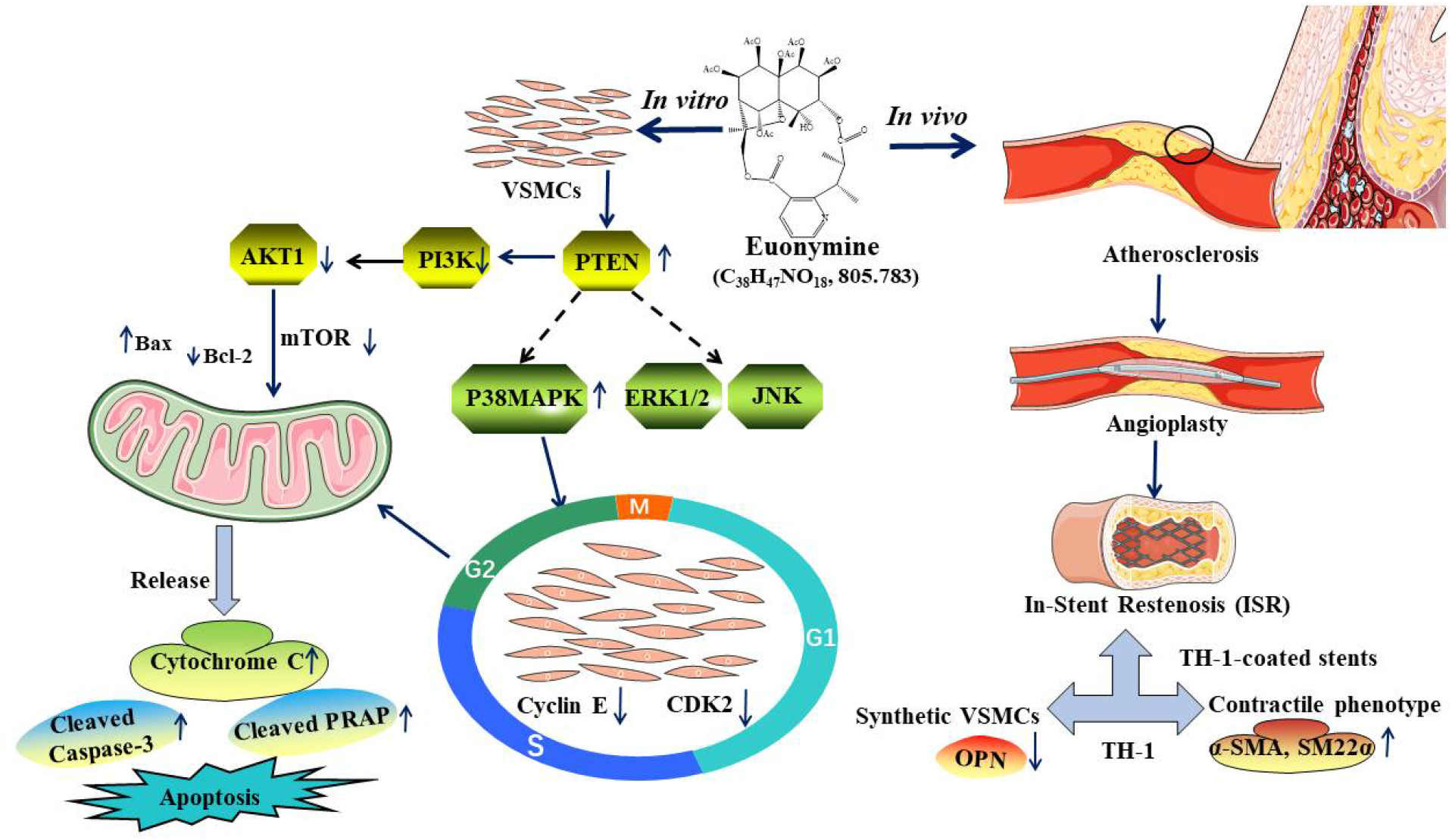

## 1. Introduction

Percutaneous coronary intervention (PCI) is the most widely used myocardial reperfusion therapeutic worldwide, which can effectively increase the survival rate and improve the quality of life of patients with coronary artery disease and myocardial infarction (1–3). However, in-stent restenosis (ISR) is a grave issue that has remained a considerable challenge in vascular surgery due to the gradual thickening of the new intima within the stent and the evolution of unstable atherosclerotic (AS) plaques after PCI(4). The development of vascular intima hyperplasia (VIH) is a common pathological feature of vascular proliferative diseases, such as atherosclerosis and ISR, which can lead to narrowing or occlusion of the lumen of blood vessels. ISR encompasses a multifactorial, multi-growth factor-mediated pathological process that involves the proliferation, migration, and dedifferentiation of multiple cells such as vascular smooth muscle cells (VSMCS) (5)(6). Exacerbated proliferation, migration, and insufficient apoptosis of VSMCs constitute the main causes of neointima hyperplasia development (7, 8). In healthy mature vessel walls, VSMCs exhibit a contractile phenotype; when the vessel is subjected to injury, VSMCs undergo a remarkable phenotypic shift from a differentiated (“contractile”) phenotype to a dedifferentiated (“synthetic”) phenotype and exhibit different morphological and functional phenotypes(9, 10). It has been well established that the synthetic phenotype of VSMCs is responsible for the occurrence of vascular proliferation and subsequent development of plastic diseases such as atherosclerosis (11), intimal hyperplasia and ISR (12, 13). Therefore, targeting and regulation of VSMC phenotypic transition and inhibition of their abnormal proliferation are important strategies considered for the prevention and treatment of intima hyperplasia. To the best of our knowledge, the effect of Euonymine on VSMC phenotypic regulation in ISR is unclear.

Previous studies have demonstrated that the phosphatidylinositol 3-kinase (PI3Κ)/protein kinase B (AKT) signaling pathway affects intracellular signaling convergence points for many factors involved in the proliferation, migration, and apoptosis of VSMCs, and plays an important role in the maintenance and regulation of cardiovascular homeostasis(14, 15). AKT, recognized as a signaling transduction and key downstream target protein of the PTEN/AKT signaling pathway, is a serine/threonine protein kinase with a wide range of substrates (16). There are three isoforms of AKT (AKT1, AKT2, and AKT3) that have been hypothesized to demonstrate distinct physiological functions (17). AKT1 is necessary for the proliferation and migration of VSMCs, and the absence of AKT1 reportedly reduces VSMC migration and survival (15). Notably, the AKT1/mTOR signaling pathway is involved in the phenotypic transformation of human and mouse aortic VSMCs and intimal proliferation in mice (18). Thus, blockade of the induction of AKT1 will be a potential therapeutic approach for ameliorating vascular occlusive diseases.

*Tripterygium hypoglaucum (Levl) Hutch*(THH) is a traditional Chinese herb that belongs to the genus Celastraceae (19). THH demonstrates an excellent immunosuppressive ability. It has also been reported that THH exhibits properties such as those pertaining to anti-inflammation, anti-tumor, and anti-fertility activities, among others (20). The chemical composition of THH is extremely complex, and approximately 300 types of components have been isolated, namely diterpenes or triterpenes such as triptolide, Celastrol, alkaloids such as sesquiterpene alkaloids, Euonymine, and so on (21, 22). Euonymine (C_38_H_47_NO_18_, 805.783) is a monomer component extracted from THH root wood and has been identified as one of the major components responsible for the exertion of immunosuppressive and anti-tumor effects of this herb.

Herein, our study demonstrated that Euonymine drug-eluting stents inhibited ISR by targeting AKT1 and p38MAPK to enhance the contractile phenotype of VSMCs to prevent intimal hyperplasia development, through the use of ox-LDL-induced cell injury model, rabbit carotid balloon injury experiments, and porcine atherosclerotic coronary implantation model. Moreover, our study provides insights into a potential therapeutic strategy involving the beneficial effect of Euonymine drug-eluting stent on ISR.

## 2. Results

### 2.1 Hematoxylin and eosin (HE) staining and immunohistochemical analysis

#### 2.1.1 Effects of Euonymine on pathological changes of intima injury in rabbit following balloon injury

On day 14 after the infliction of balloon injury on the intima, the histomorphology of the vessels was investigated. (1) HE staining revealed that the sham-operated group presented with uniform wall thickness, marked layers, intact internal elastic plates, and the absence of VSMCs in the intima; the model group demonstrated the presence of pathological thickening of the intima to a certain extent, along with disorganized VSMCs, and exhibited migration and proliferation from the media to the intima; the Euonymine (0.25 mg/mL) group demonstrated the development of mild neoplastic intimal hyperplasia, along with the presence of disorganized VSMCs in the neoplastic endothelium, fibrous tissue, and less extent of luminal stenosis, while Euonymine (0.5 mg/mL) group exhibited the presence of ordered VSMCs in the vascular smooth muscle intima, less extent of neoplastic intima, and did not present with significant luminal stenosis (Figure 2. A-a). (2)Vascular histomorphometric analysis showed that the lumen area was reduced by 37.3% in the model group compared to the sham-operated group, and by 25.7% and 11.0% in the Euonymine group (0.25 mg/mL, 0.5 mg/mL), respectively; additionally, compared to the model group, the neonatal intimal area was reduced by 62.5% and 86.9% in the Euonymine group (0.25 mg/mL, 0.5 mg/mL), respectively (P<0.01), and the N/M ratio was reduced by 57.6% and 81.4% (P<0.01), respectively, as shown in Figure 2. A-b.

**Figure 1.**
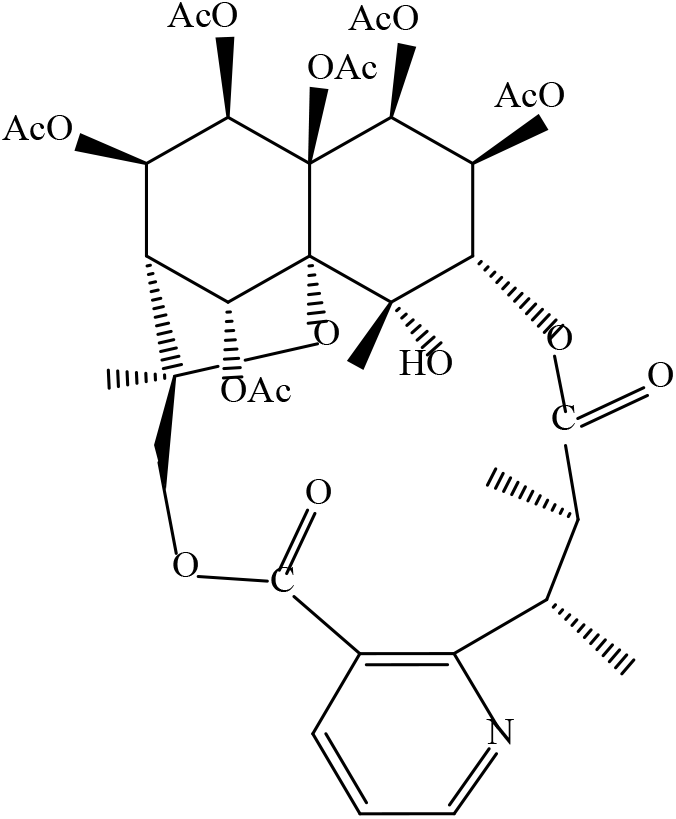
Chemical structure of Euonymine.

**Figure 2.**
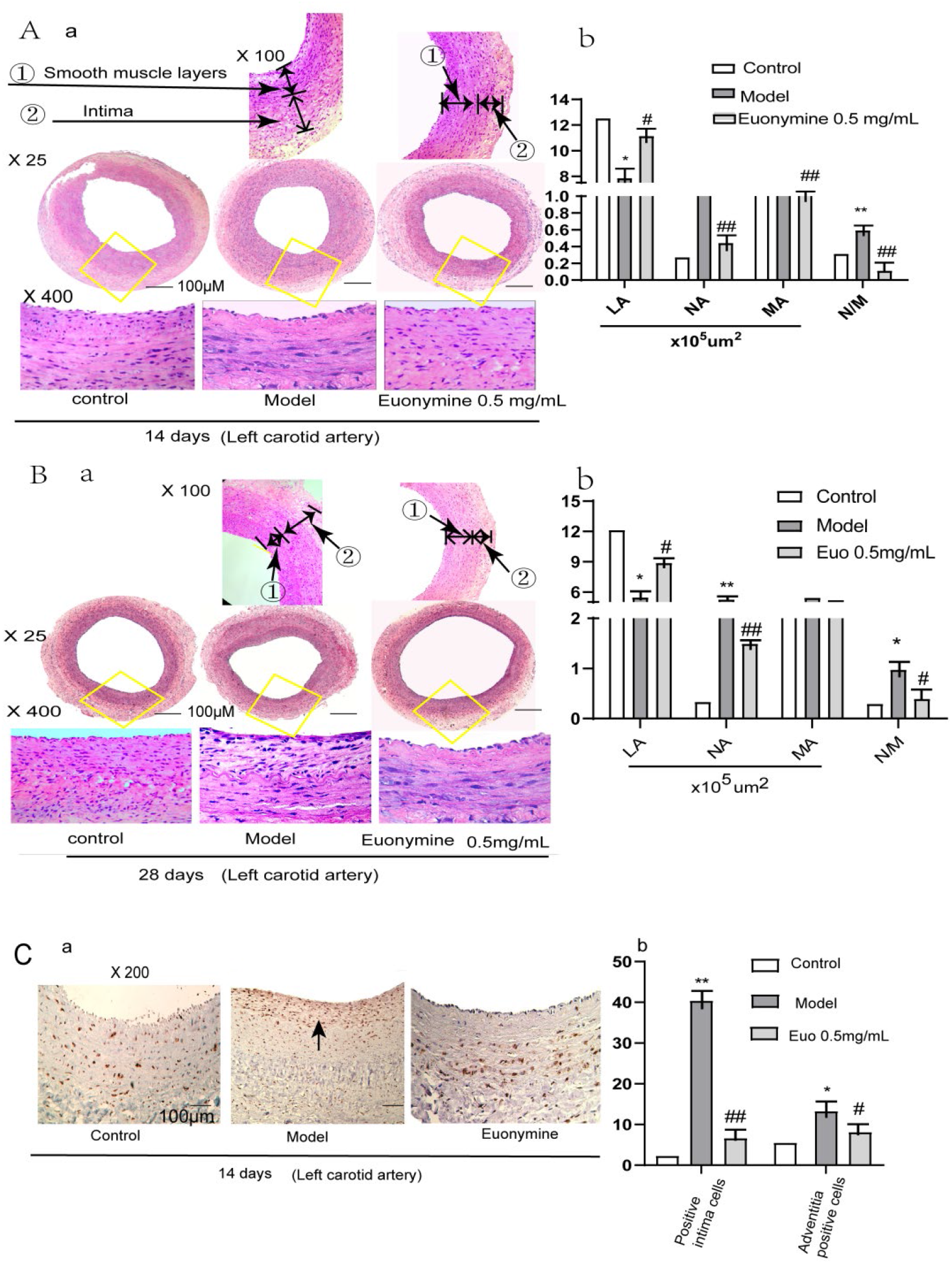
Euonymine attenuates neointimal hyperplasia in rabbit carotid arteries following balloon injury. A and B, Euonymine attenuates neointimal hyperplasia in rabbit carotid arteries following balloon injury. A,(A-a and B-a), Representative hematoxylin and eosin (HE)–stained carotid artery sections derived from rabbit treated with Euonymine (0.5mg/mL) following balloon injury on days 14 and 28 after infliction of injury. (HE staining; top, ×100, ×25 magnification; bottom, ×400 magnification). B (A-b and B-b), The effects of Euonymine on neointimal hyperplasia are quantitated by neointimal area and intimal hyperplasia (ratio of intima/media area, N/M; *P<0.05, **P<0.01 vs. the control group; ^#^P<0.05 vs. model group). Comparison of the lumen area, intimal and medial area, N/M ratios among groups. LA, lumenal area; NA, neointimal area; MA, medial area; n=4. (C), Representative photomicrographs of PCNA–expressing arterial sections on day 14 after infliction of injury. a, (IHC, ×200 magnification).b, Analysis of the percentage of PCNA-positive cells vs. total number of VSMCs. n=5. Scale bar= 100μm.

On day 28 after the infliction of balloon injury on the intima, the histomorphology of the vessels was investigated. (2) HE staining revealed no significant changes in the sham-operated group compared to those observed in the group after 14 days; in the model group, the injured intima presented with significant thickness. The lumen was found to be comparatively small, centripetal, or eccentric, and a substantial number of VSMCs presented with migration from the media to the intima, along with proliferation. The nuclei of VSMCs in the new intima were observed to be disorganized, and were significantly different from those in the media. Compared with the model group, the Euonymine group (0.25 mg/mL, 0.5 mg/mL) demonstrated a dose-dependent increase in the lumen area and decrease in neointima formation, as shown in Figure 2 B-a. (2)Vascular histomorphometric computer image analysis revealed that the lumen area of the model group decreased by 45.6% compared to that of the sham-operated group observed on day 28 postoperatively, while the lumen area of the Euonymine group (0.25 mg/mL, 0.5 mg/mL) decreased by 39.4% and 31.6%, respectively. Compared to the model group, the lumen area of the Euonymine-administered group decreased by 41.6% and 33.5% (P < 0.01), respectively, and the N/M ratio decreased by 42.3% and 26.9% ( P < 0.01), respectively, as shown in Figure 2 B-b.

On day 28 after the infliction of balloon injury, immunohistochemical analysis was performed; immunohistochemical staining results showed that proliferating cell nuclear antigen (PCNA) expression was widely distributed in the intima and media in the model group, while only a few PCNA-positive cells were found in the arterial intima and media in the Euonymine group, as depicted in Figure 2. C-a. By analyzing the percentage of positive PCNA-stained cells in the intima and media VSMCs, 14 days after the infliction of balloon injury, the statistical results showed that the ratios of PCNA-positive cells in the neointima of the Euonymine-administered group(0.25 mg/mL, 0.5 mg/mL) were 17.68±5.27 and 6.60±2.52, and both were found to be significantly less compared with the model group presenting with a ratio of 40.37±7.12 ( P < 0.01). Additionally, the ratios of PCNA-positive cells in the media of the model and the Euonymine-administered group were 13.24±3.59, and 12.11±3.27, 10.88±1.87, respectively, as shown in Figure 2. C-b.

#### 2.1.2 Euonymine effectively inhibits neointimal hyperplasia in porcine atherosclerotic coronary implantation models

The vascular tissue boundary between the stent segment and non-stent segment was remarkable, as shown in Figure 3 A. Twenty-eight days after stent implantation, the stent vessels were subjected to staining with HE and were observed, as shown in Figure 3 B. The results showed that the neointimal thickness, neointimal area, and stenosis degree in the high dose Euonymine-coated stent group were similar to those in Sirolimus-eluting stent (SES) group; however, the values were smaller than those obtained in other experimental groups ( P < 0.05). Inflammation scores were greater in the model group than those noted in Bare-metal stent (BMS) group (P<0.01); the presence of Euonymine led to a decrease in inflammation scores in the injured intima cases compared to the model group cases. Moreover, no statistically significant difference in inflammation scores was noted in the high dose Euonymine group compared with the SES group ( P > 0.05), as shown in Figure 3. C.

**Figure 3.**
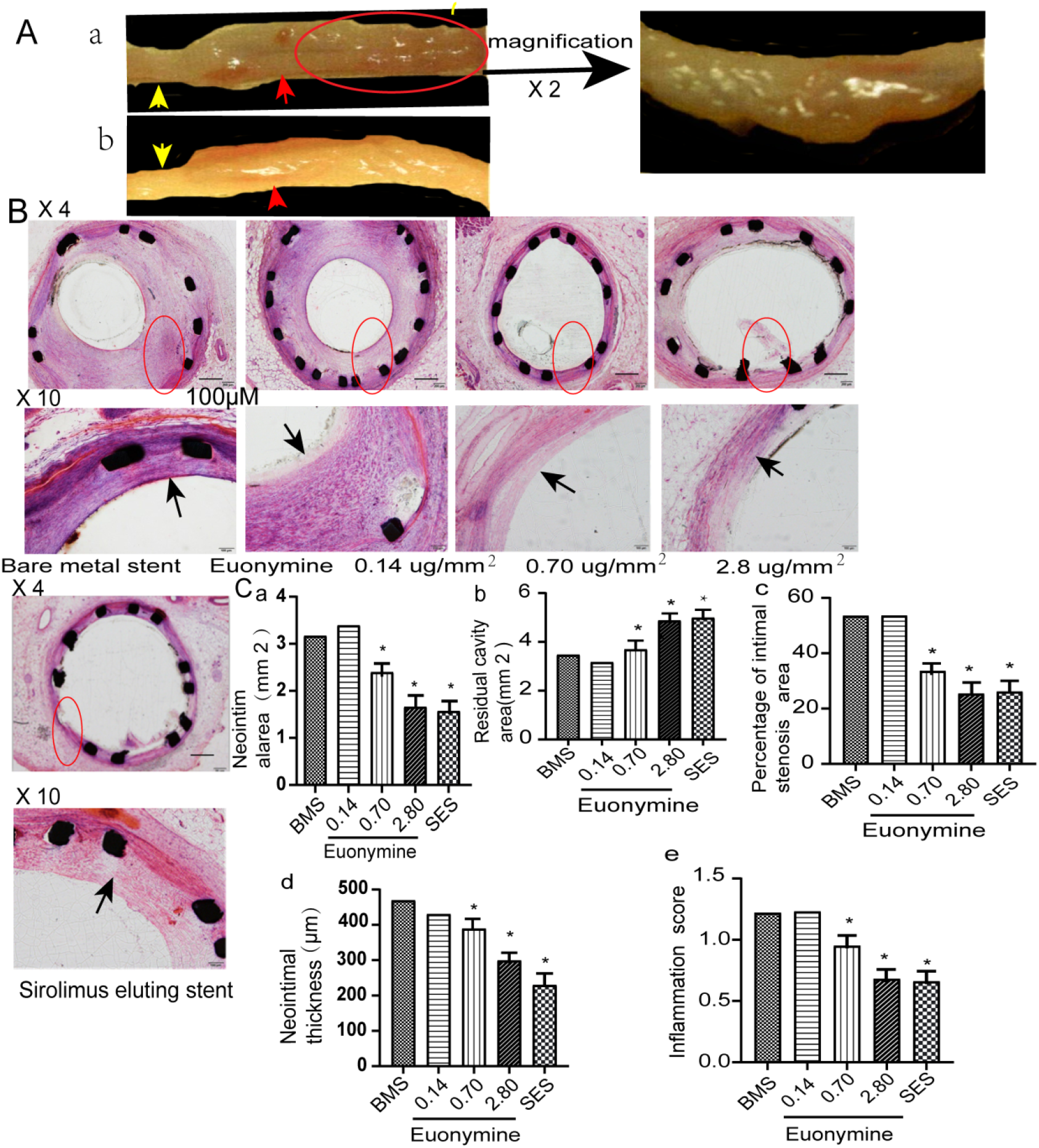
Euonymine effectively inhibits neointimal hyperplasia in porcine atherosclerotic coronary implantation models. A, (a-b), The vascular tissue boundary between the stent segment and non-stent segment is evident. The yellow arrow indicates a non-stent segment of vascular tissue, and the red arrow indicates a stent segment of vascular tissue. B, Hematoxylin and eosin (HE) staining of stent segments in each group, (HE staining; top, ×4 magnification; bottom, ×10 magnification). The black arrow indicates the recruitment of inflammatory cells to the vascular tissue. C, Morphological parameters of vascular restenosis in implanted segments. Bare-metal stent (BMS), three Euonymine treatment groups ( 0.14 [low-dose group], 0.70 [medium-dose group], and 2.80 [high-dose group] μg/mm^2^), and Sirolimus-eluting stent (SES). *P<0.05, vs. the BMS group. Data have been presented as mean ± SD (n = 4). Scale bar =100μm.

### 2.2 Euonymine effectively reduces vascular stenosis on porcine coronary arteries after stent **implantation.**

The coronary angiography results obtained on day 28 after stent implantation showed no significant luminal stenosis in the high-dose and SES groups, while the BMS, low-, and medium-dose groups showed varying degrees of luminal stenosis, as shown in Figure 4. Our experimental results suggested that the medium- and high-dose groups of Euonymine-coated stents presented with effective inhibition of coronary artery neointimal hyperplasia in the porcine coronary artery model, while the low dose group presented with no significant effect. Qualitative comparative analysis (QCA) has been considered an important method and gold standard for the diagnosis of coronary artery disease and for the evaluation of coronary stenosis, since its clinical use in 1958; however, it presents with limitations as it can only be performed to reflect the lumen of the vessel and cannot reflect wall-related changes (23, 24). Owing to the limitations of angiography, we performed optical coherence tomography (OCT) for the injury model of neointimal hyperplasia.

**Figure 4.**
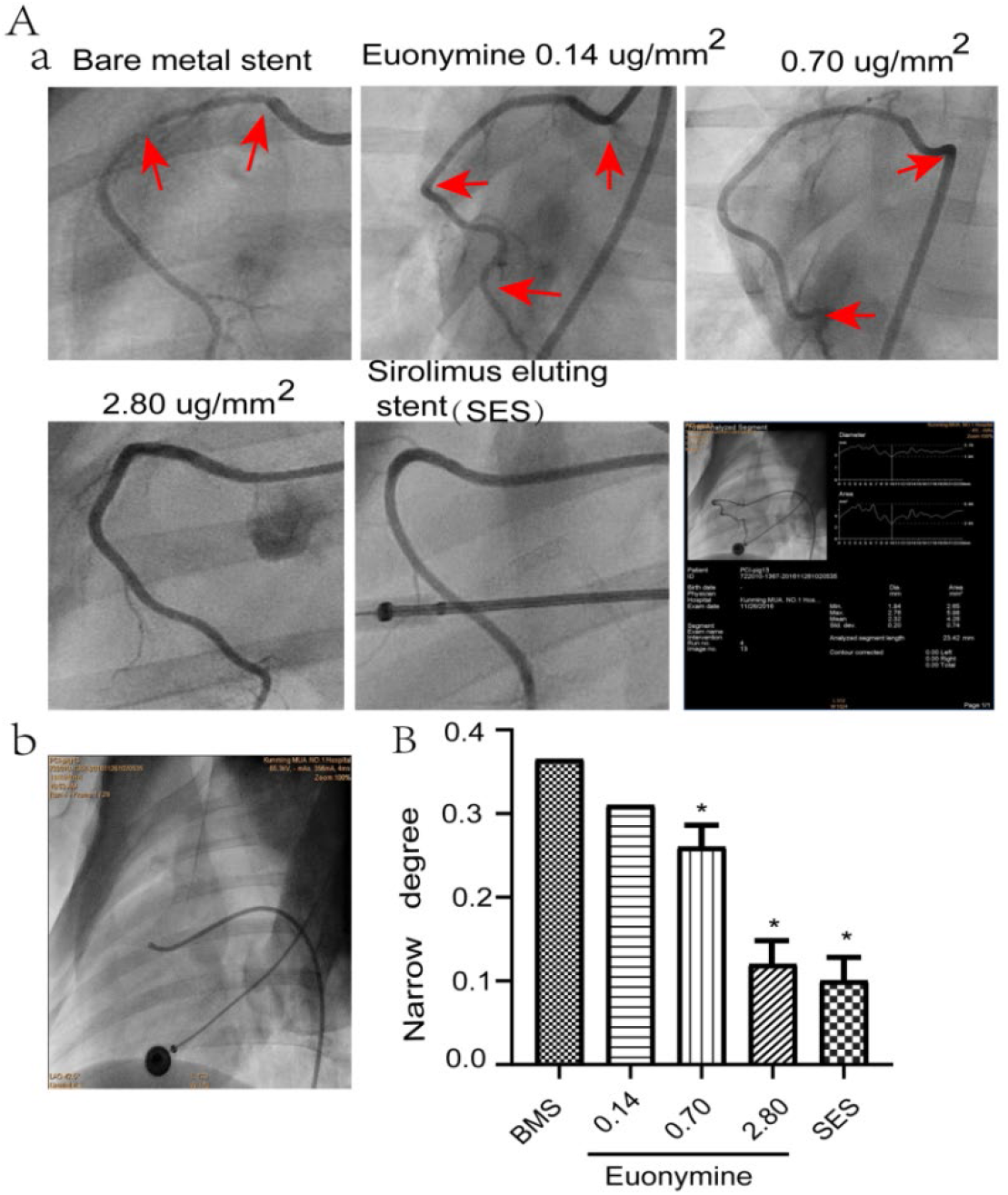
Coronary angiography after stent implantation on day 28. A, (a and b), a represents angiographic results on day 28 after stent implantation in different groups. b represents on immediate post-procedural angiogram, angiography shows that the stent was dilated completely. The red arrow indicates the narrowing of coronary artery. *P<0.05, **P<0.01 vs. the Bare mental stent group (BMS). Means ± SD, (n = 3); B, Results of coronary angiography and quantitative analysis.

### 2.3 Euonymine effectively suppresses ISR after PCI on porcine coronary arteries

OCT is an *in vivo* examination that can help compensate for the shortcomings of QCA and has garnered attention as the new gold standard for intravascular imaging of stents, atherosclerotic progression, vulnerable plaques, and neointimal hyperplasia (25, 26). The neointimal hyperplasia proliferation of each stent was observed continuously and throughout the procedure via OCT. The results of OCT suggested that compared to the negative control group, the neointimal thickness and the extent of stenosis were reduced in the medium- and high-dose Euonymine-coated stent groups (P < 0.05), while the residual lumen area values in both the high-dose Euonymine-coated stent group and the positive control group were significantly greater than those noted in other experimental groups (P < 0.05). Furthermore, different degrees of neointimal hyperplasia and lumen shrinkage were observed in the model group and Euonymine low- and medium-dose groups, while no significant neointimal hyperplasia was observed in the high-dose Euonymine-eluting stent group and Sirolimus-eluting stent group. As shown in Figure 5. Furthermore, the results of OCT were consistent with those obtained after HE staining and coronary angiography.

**Figure 5.**
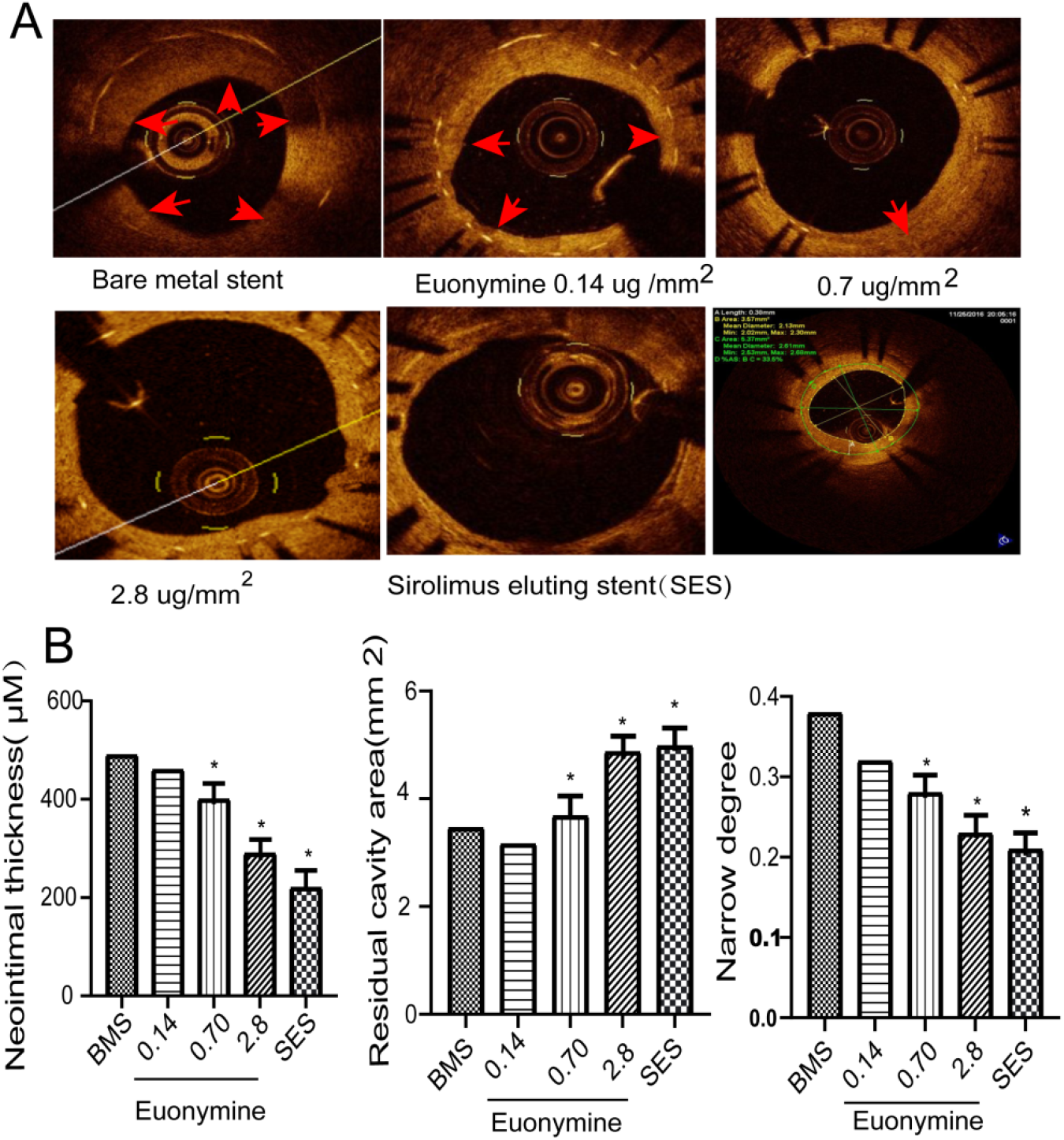
OCT Images of porcine coronary arteries after stent implantation on day 28. A, representation of *in vivo* OCT results obtained for different stent groups. Different degrees of intimal hyperplasia and lumen shrinkage could beobserved in the Bare-metal stent group (BMS), low-dose, and medium-dose Euonymine-eluting stent groups. However, no significant intimal hyperplasia is found in the high-dose Euonymine-eluting stent group and Sirolimus-eluting stent group. Red arrows indicate significant intimal hyperplasia. B, Quantitative analysis results obtained for OCT measurement. *P<0.05, **P<0.01 vs. the SES group. Data have been presented as mean ± SD (n = 4).

### 2.4 Euonymine attenuated neointimal hyperplasia via inhibition of phenotypic transition of VSMCs and regulation of the PTEN/AKT1/mTOR pathway in rabbit following balloon injury

To further investigate the phenotypic transformation of VSMCs in neointimal hyperplasia, we performed immune fluorescence (IF) and western blotting (WB) analyses to label contractile phenotypic markers [anti-α-smooth muscle act in (α-SMA) and smooth muscle protein 22α (SM22α)] and synthetic markers (osteopontin) in rabbit following balloon injury. Based on IF in vascular tissue revealed that the fluorescence intensity of SM22α and α-SMA was decreased in the model group, while the expression of osteopontin was increased. Interestingly, we found that Euonymine attenuated the phenomenon in a dose-dependent manner, as shown in Figure 6. B. Furthermore, the results were confirmed in parallel in vascular tissue via WB, as shown in Figure 6. A.

**Figure 6.**
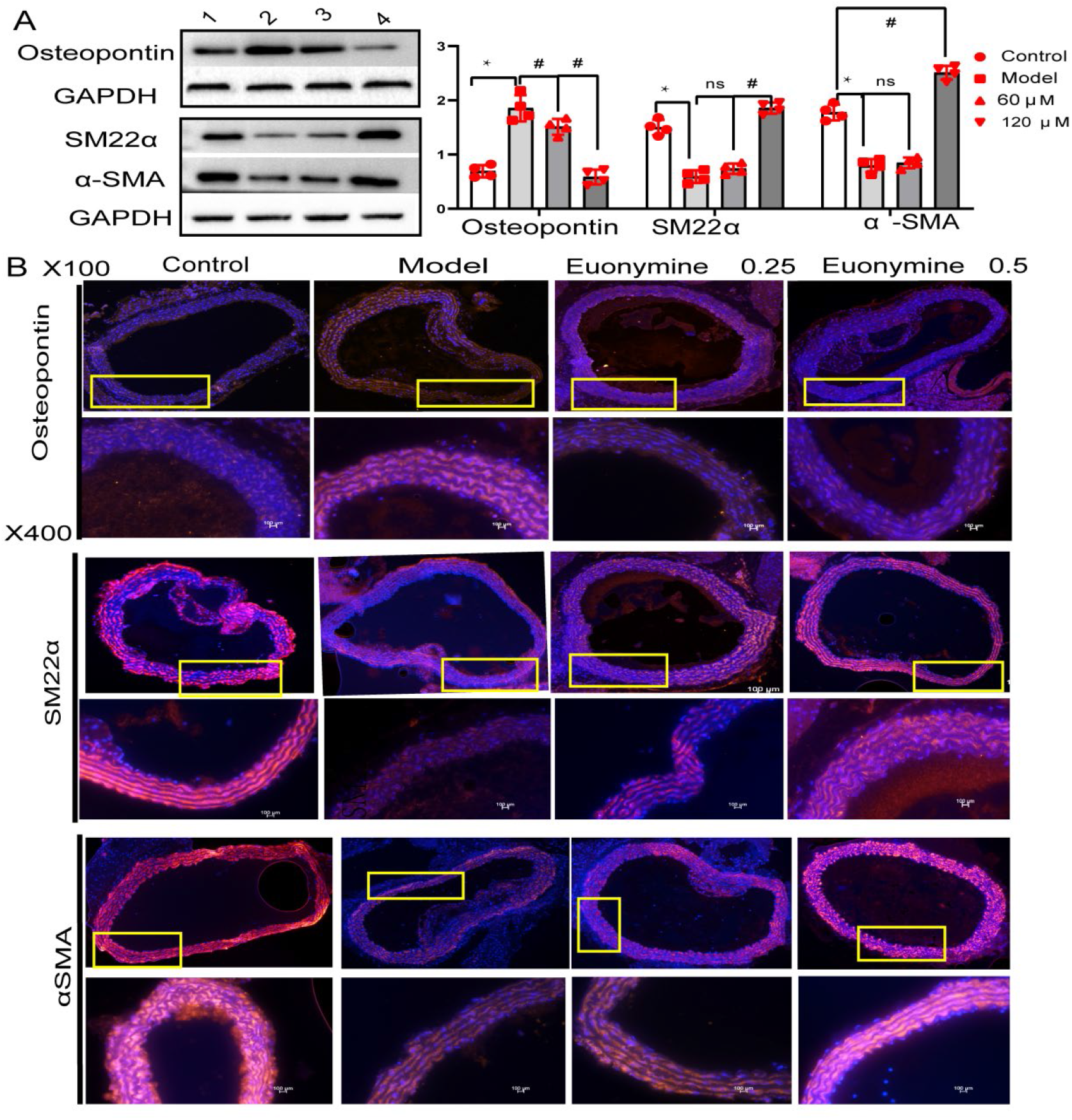
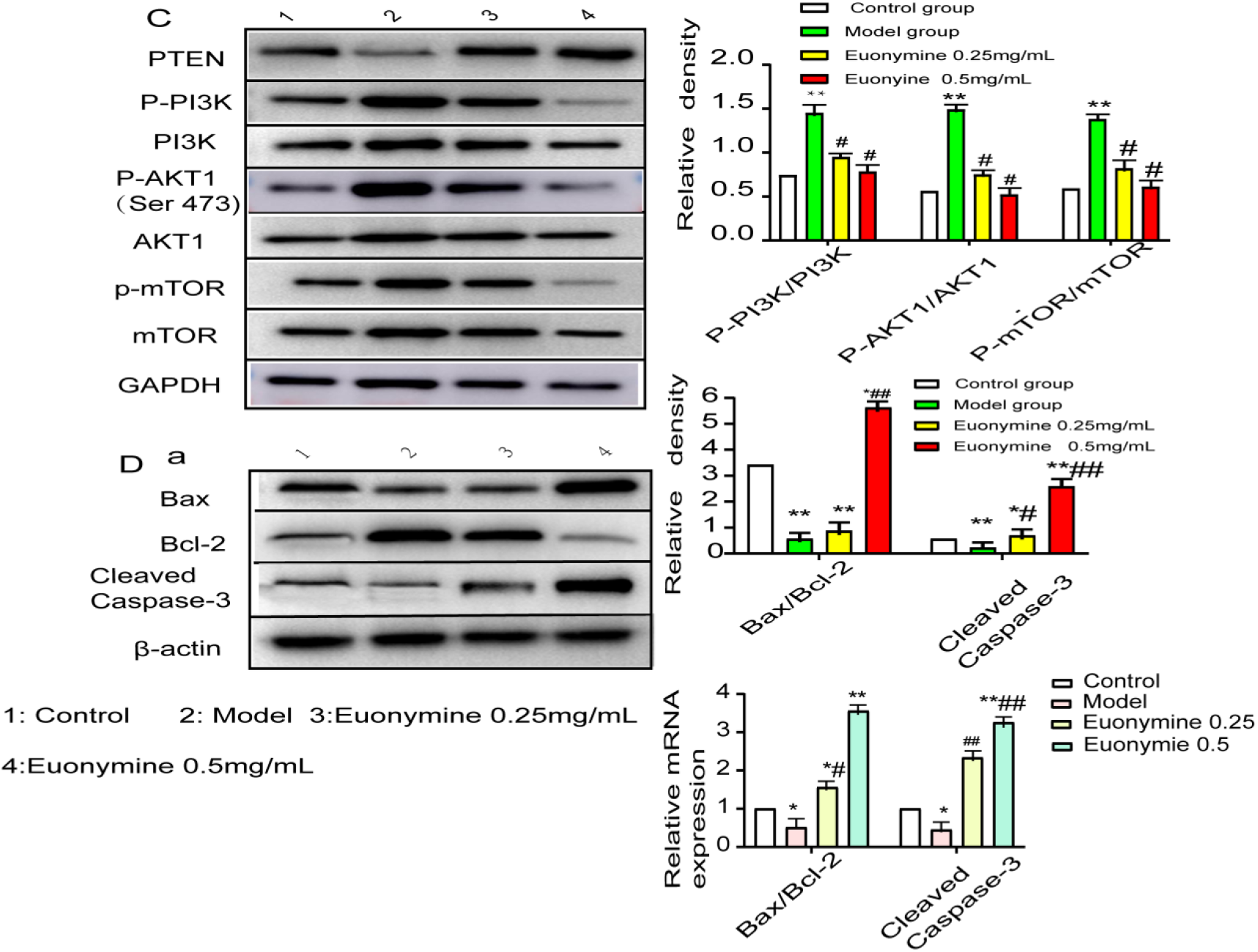
Euonymine attenuated intimal hyperplasia via inhibition of phenotypic transition of VSMCs and mediation of PTEN/AKT1/mTOR in rabbit following balloon injury. A, Protein levels of osteopontin, SM22α, and α-SMA were determined via western blotting (WB) after Euonymine treatment for 14 days. B, Immunostaining analysis performed for osteopontin, SM22α, and α-SMA expression; scale bar = 100μm. C, The phosphorylation levels of the PTEN/AKT1/mTOR pathway-related proteins and total protein content measured via WB. D, The apoptotic signal signaling pathway-related proteins were evaluated via WB and RT-PCR. The relative protein expression levels have been normalized to those of GAPDH or β-actin and have been quantified using the ImageJ v1.49 software. Data have been presented as mean ± SD. *P< 0.05, and ** P< 0.01 vs. the control group; ^#^P < 0.05 and ^##^P < 0.01 vs. the model group; n=6.

To explore the possible effects of Euonymine on neointimal hyperplasia through the PTEN/AKT/mTOR signal pathway, WB and reverse transcription polymerase chain reaction (RT-PCR) were performed. WB and RT-PCR results showed that the model group presented with a significant increase in the phosphorylation levels of PI3K, AKT1(Ser473), and mTOR, compared with the control group. Additionally, the use of Euonymine significantly decreased the phosphorylation levels of PI3K, AKT1, and mTOR in the model group. Moreover, as a negative regulatory molecule of P13K, the expression trend of PTEN helped verify the above-mentioned experimental results, as shown in Figure 6. C. Furthermore, we also determined whether Euonymine affected apoptotic signals in rabbits following angioplasty. Angioplasty caused upregulation of Bcl-2 expression and downregulation of expression of Bax and cleaved caspase-3, which were attenuated by Euonymine. Bax/Bcl-2 ratio was evaluated to reflect the apoptosis level, and the results indicated that cell apoptosis was inhibited by balloon injuries. However, cell apoptosis was induced by Euonymine.

Taken together, it may be preliminarily suggested that Euonymine may inhibit neointimal hyperplasia development in models with rabbit balloon injury to carotid artery, partly by inducing a VSMC contractile phenotype that can present with modulation of the PTEN/AKT1/mTOR pathway.

### 2.5 Euonymine mediates phenotypic switching of VSMC via AKT1 to suppress VSMC proliferation and migration

To explore the mechanism of Euonymine-mediated inhibition of intimal hyperplasia development, the effects of Euonymine on the proliferation of aortic VSMCs *in vitro* were analyzed and observed. The growth of aortic VSMCs was monitored via phase-contrast microscopic morphological observation and anti-α-actin monoclonal antibody staining, as shown in Figure 7A. To investigate the cytotoxic effect of Euonymine on VSMCs, cells were subjected to exposure to increasing concentrations of Euonymine for five time points; cell viability was subsequently evaluated via 3-(4, 5)-dimethylthiazol-z-yl)-3,5-di-phenyltetrazoliummromide (MTT) assay and cell viability, as shown in Figure 7. B and C. Euonymine significantly reduced the viability of VSMCs in both dose-dependent and time-dependent manner. Furthermore, we found that Euonymine-treated VSMCs exhibited a typical spindle-like or triangular intact appearance, tending towards a contractile type with a reduction in cell volume and number, as shown in Figure 7. E. The median inhibitory concentration (IC_50_) of Euonymine for VSMCs was estimated to be 33.92 μg/mL at 48 h, as shown in Figure 7 D.

**Figure 7.**
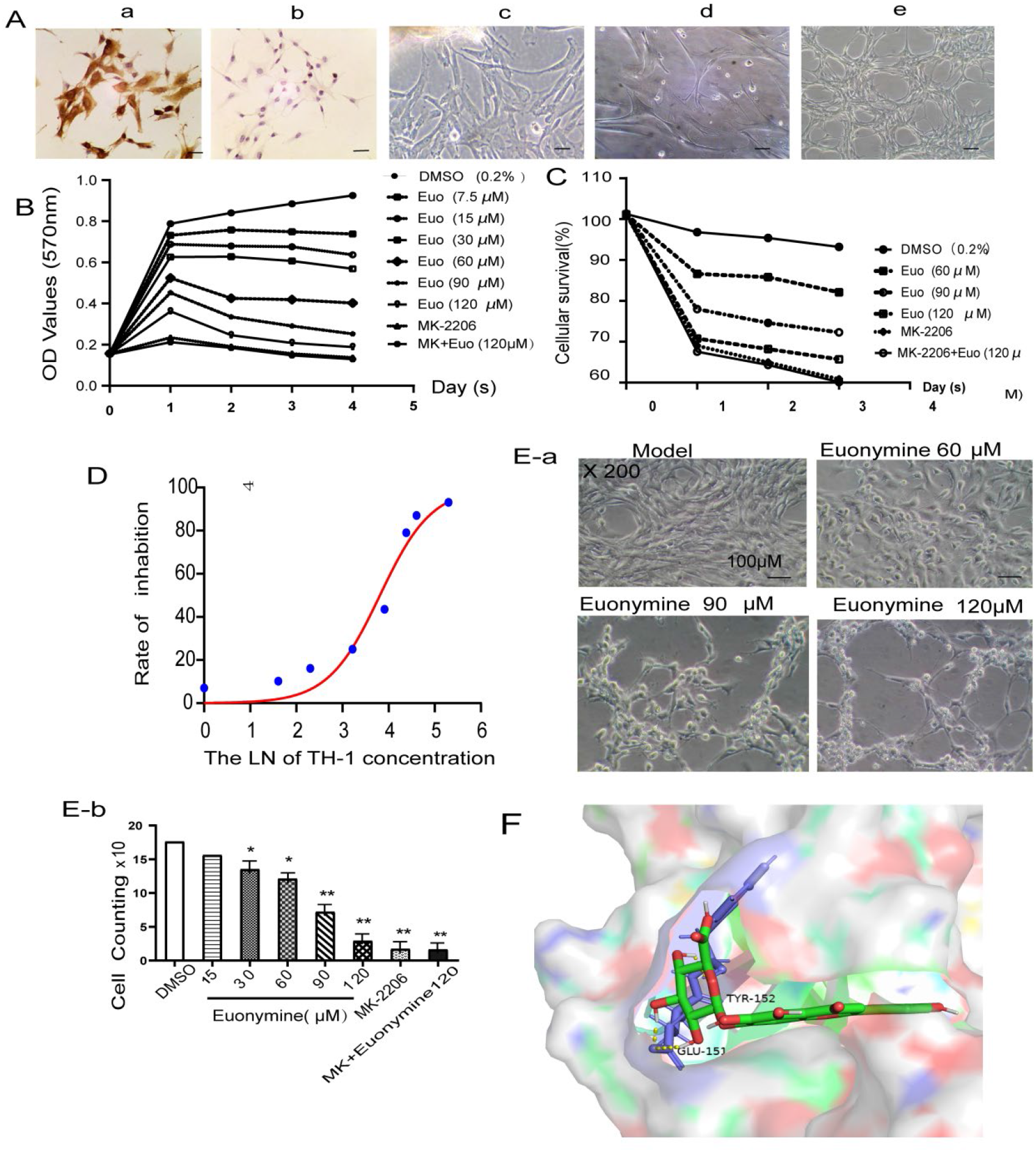
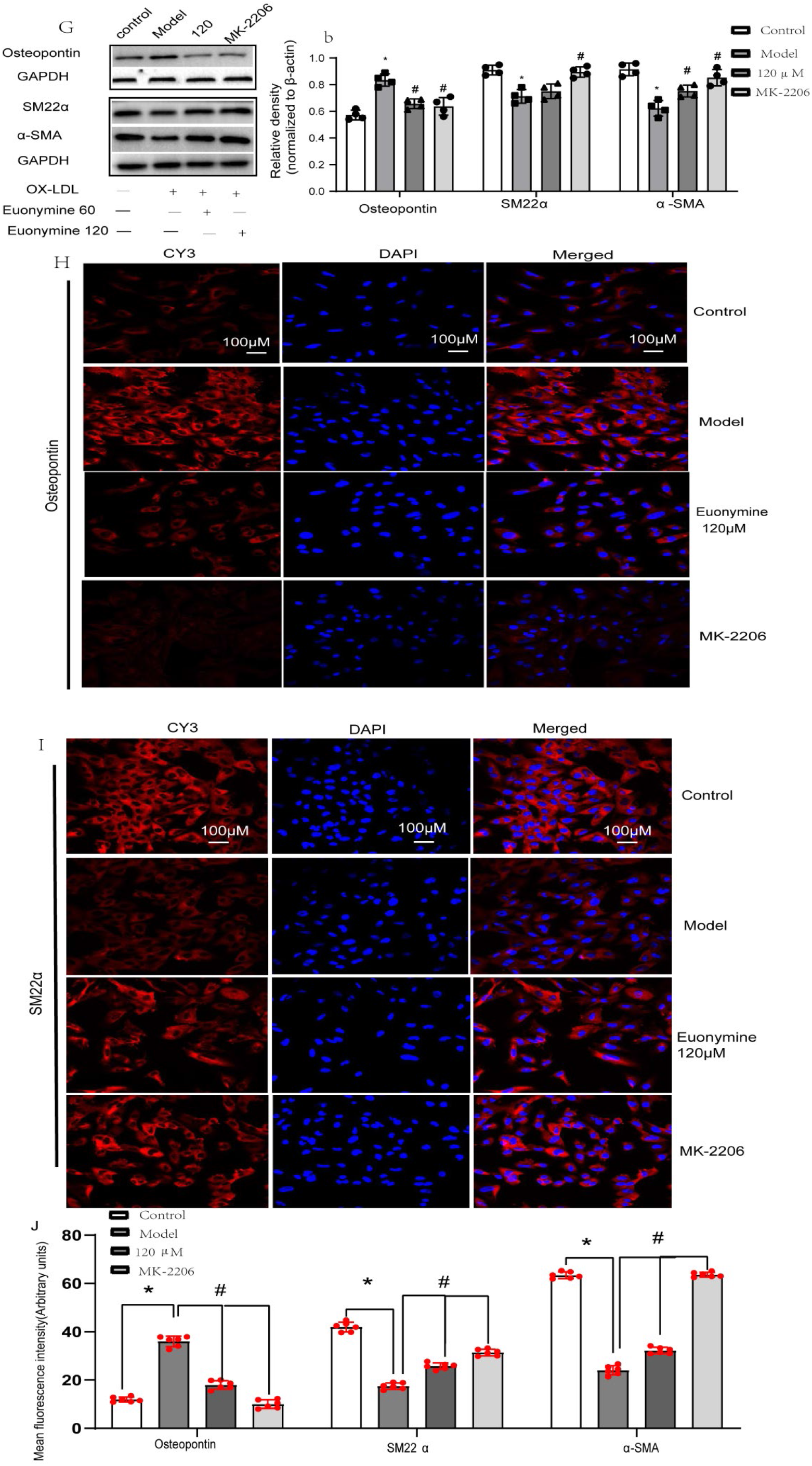
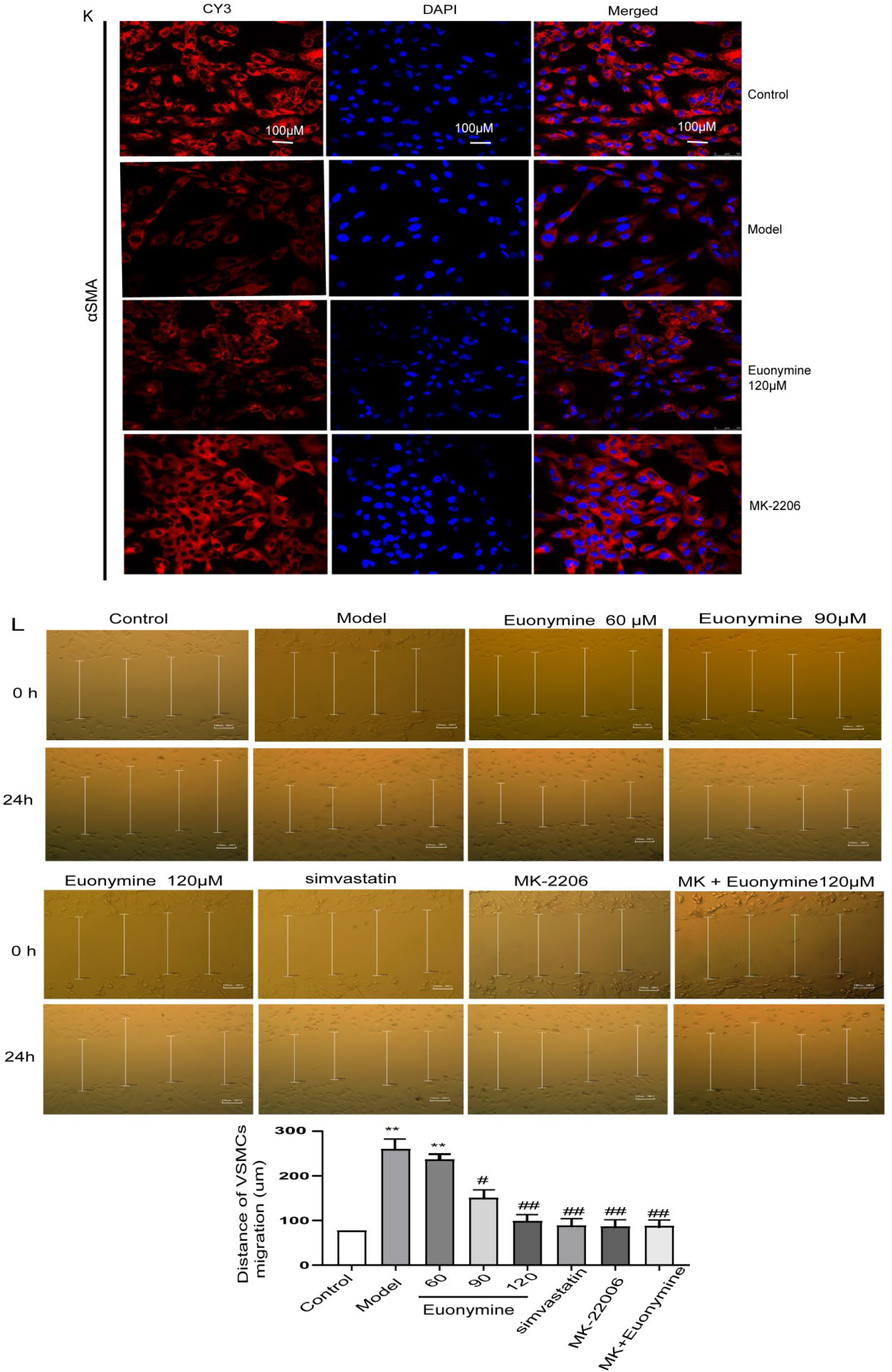
Euonymine mediates phenotypic switching of VSMCs via AKT1 to suppress VSMCs proliferation and migration. A, Typical appearance of VSMCs under alight microscope. (a, b) Identification of VSMCs via immunostaining with α α-actin antibodies. a. Positive cells. b. Negative control. (c, d) Primary cultured VSMCs (×200 magnification). e, VSMCs obtained after 6 passages. (×100 magnification). B, Dependent curve of concentration and time depicted for VSMC inhibited by Euonymine (demonstrated via MTT assay). C, Euonymine inhibits cell viability. VSMCs were incubated for indicated periods with Euonymine (60, 90, and 120µM). Viable cells have been counted after staining with trypan blue. Data represent the average of three experiments. D, Curve for inhibition rate noted after Euonymine treatment. E, (a and b), a reprents decrease in the number of VSMCs. We found that Euonymine-treated VSMCs exhibited a typical spindle-like or triangular intact appearance, tending towards a contractile type with a reduction in cell volume and number. b reprents the effects of different concentrations of Euonymine on the proliferation of VSMCs estimated by using the Cell Counting. F, Diagram of the molecular docking pattern of AKT1. Molecular docking results showed good docking activity of Euonymine and AKT1 targets. Purple represents AKT1 docking sites, green represents Euonymine; the numbers in the b diagram represent amino acid residues. Binding energy(kcal/mol) is −2.25. Finally, the AutoDock4.2.6 software was used to verify the molecular docking between the selected chemical component and the core target spots. G, Protein levels of osteopontin (OPN), SM22α and α-SMA determined via western blotting(WB) after Euonymine treatment for 24h. H-K, Immunostaining for osteopontin, SM22α, and α-SMA expression; scale bar = 100μm. L, Euonymine inhibits migration of VSMCs. Serum-starved VSMCs were subjected to stimulation with 75 mg/mL ox-LDL in the presence or absence of Euonymine (concentration range of 60-120µM), or simvastatin (10µM), or MK-2206 (30µM) for 24 h. a. Representative images showing migration of VSMCs at 0 h and 24 h (×100 magnification). b. Quantitative data have been presented as the percentage of VSMCs migrating into the wound with respect to the cell-free area at 0 h; n=6. *P<0.05 vs. 0.2% DMSO (control), ^#^P<0.05 vs. ox-LDL (75mg/mL). Scale bar = 150μm.

We further validated the regulation of phenotypic transition of VSMCs via Euonymine exposure at the cellular level. To further confirm the mechanism of drug inhibition of VSMCs proliferation, we used molecular docking technique to identify the Euonymine targets of action. Molecular docking results showed good docking activity of Euonymine and AKT1 targets, as shown in Figure7. F. Next, we performed IF and WB analyses to label contractile phenotypic and synthetic markers. Based on IF analysis, it was inferred that the fluorescence intensity of SM22α and α-SMA increased, while osteopontin expression decreased in Euonymine and MK-2206 (a highly selective inhibitor of AKT1; it inhibits the auto-phosphorylation of AKT1 Thr308 and Ser473) groups, compared with the ox-LDL-induced group, but no difference between Euonymine and MK-2206 group was noted, as shown in Figure 7.G-K. The results were confirmed in parallel in vascular tissue via WB, as shown in Figure 6. A and B. Taken together, these findings revealed that AKT1 mediated VSMC phenotypic switching. Moreover, wound healing-induced migration assay showed that the ox-LDL-injured cell migration was inhibited by Euonymine in a dose-dependent manner, as shown in Figure 7. L.

In summary, our data indicate that Euonymine potently mediates the proliferation and migration of VSMCs during neointimal hyperplasia via AKT1 inhibition of synthetic phenotype of VSMCs. Additionally, no cytotoxicity was noted.

### 2.6 Euonymine regulates the PTEN/AKT1 signaling pathway via inhibition of the AKT1 phosphorylation site

To further confirm that Euonymine treatment was affected by the PTEN/AKT1 signaling pathway *in vitro*, we performed WB, IF, and RT-PCR analyses, as shown in Figure 8. Our data showed that phosphorylation levels of AKT1 were significantly reduced after 24 h of Euonymine treatment of VSMCs. In contrast to the decrease noted in phosphorylated AKT1 expression, PTEN levels were found to be significantly increased after Euonymine treatment in a dose-dependent manner. Noticeably, ox-LDL leads to the activation of the AKT1 (Ser473) and AKT1 (Thr308) sites, an activity found to be reversed via Euonymine treatment. To further confirm the role of AKT1 activation in Euonymine-induced apoptosis, VSMCs were subjected to pre-treatment with AKT1 inhibitor MK-2206 (30μM) prior to Euonymine treatment. The ox-LDL-induced activation of PTEN/AKT1 was almost eliminated by MK-2206 pre-treatment. The above-mentioned results suggested that Euonymine regulated the PTEN/AKT1 signaling pathway by inhibiting the expression of the phosphorylation site AKT1, thereby inhibiting ox-LDL-induced proliferation and migration.

**Figure 8.**
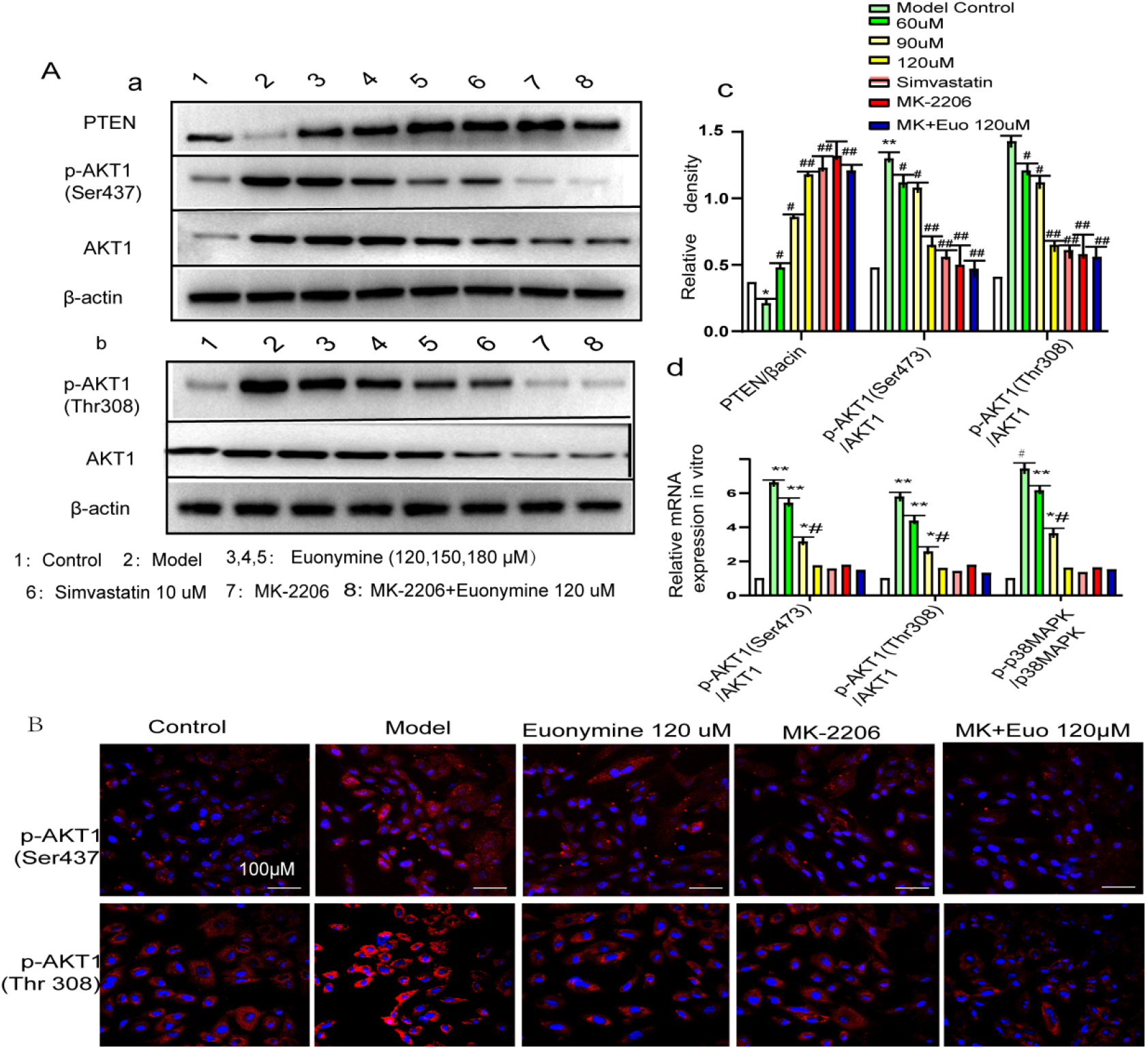
Euonymine regulates the PTEN/AKT1 signaling pathway via inhibition of the AKT1 phosphorylation site. Serum-starved VSMCs were subjected to pre-treatment with Euonymine (60μM, 90μM, 120μM) for 24 h, followed by treatment with 75mg/L ox-LDL for 24h. Thereafter, VSMCs were subjected to treatment with MK-2206 30uM in the absence or presence of Euonymine (120μM) for 24h. A, The expression levels of PTEN, p-AKT1(Ser437), p-AKT1(Thr308) have been determined via western blotting (WB) and RT-PCR. B, Representation of immunofluorescence results of p-AKT1(Ser437) and p-AKT1(Thr308) (×200 magnification). The bands of phosphorylated proteins have been normalized to those of total protein expression or GAPDH/β-actin. Data have been presented as mean ± SD. *P< 0.05, and ** P<0.001 vs. the control group; ^#^P < 0.05 and ^##^P < 0.01 vs. the ox-LDL group; n=6 per group. Scale bar=100μm.

### 2.7 Euonymine triggers apoptosis of VSMCs *in vitro*

To further explore the mechanism of anti-neointimal hyperplasia via Euonymine treatment, we investigated the apoptosis-inducing effect of Euonymine on VSMCs by adopting various methods. First, we observed the morphological changes of VSMCs through Giemsa staining, and our data demonstrated the occurrence of morphological changes in early apoptosis and cell necrosis induced by different concentrations of Euonymine in VSMCs, as shown in Figure 9 A. Subsequently, the TUNEL labeling assay revealed that few nuclei possessed brown-yellow particles and few nuclei were pyknotic or fragmented into several brown-yellow particles (indicated using a yellow arrow) in Euonymine-treated group; additionally, the nuclei did not exhibit staining or only a few brown-yellow nuclei were observed in the control group (0.2% DMSO), as shown in Figure 9. B. Furthermore, intracellular morphological changes of VSMCs were further examined via transmission electron microscopy (TEM). TEM analysis showed that typical apoptotic changes and apoptotic bodies were observed in VSMCs subjected to treatment with Euonymine, as shown in Figure 9. C. Moreover, our data suggested that 8.1% of the apoptotic cells and 0.3% of the secondary necrotic cells could be observed in the TUNEL/PI double-labeled parameter diagram. With the increase of Euonymine concentration, the results of flow cytometry analysis showed that the number of apoptotic cells (indicated in the green fluorescence region) increased significantly with the increase of Euonymine concentration and presented with values up to 39.3%. The secondary necrotic cell proportion ( indicated in the red fluorescence region) increased slightly, however, no significant difference was noted between low and high concentration groups (P>0.05), as shown in Figure 9. D. These results confirmed that the inhibitory effect of Euonymine on VSMC proliferation was mainly executed through apoptosis induction rather than direct cytotoxicity, as shown in Figure 9. E.

**Figure 9.**
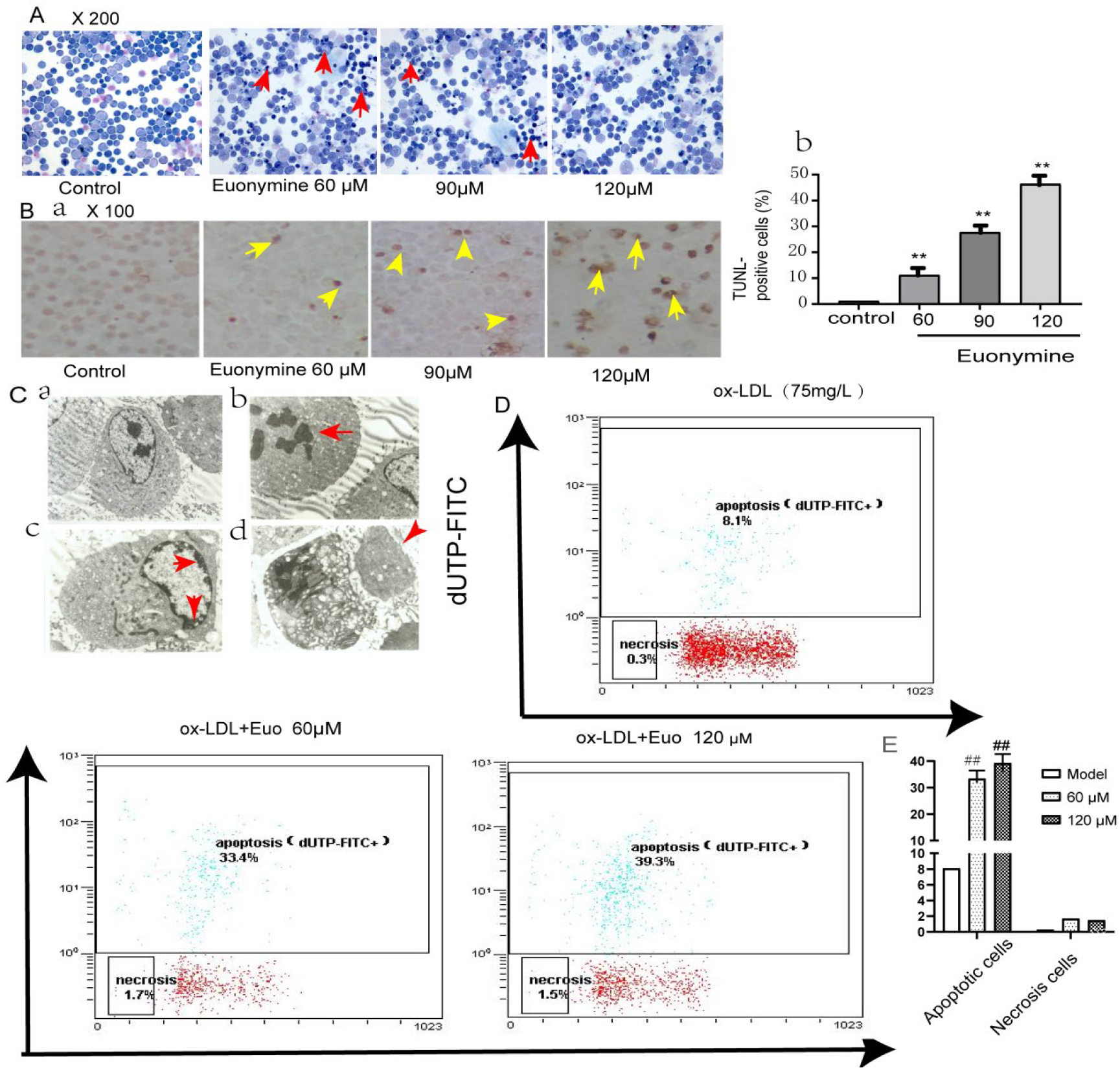
Euonymine induces apoptosis in VSMCs. A, Representation of morphological and nuclear changes of apoptotic VSMCs induced by Euonymine treatment (Giemsa staining, ×200 magnification). Serum-starved VSMCs were subjected to treatment with 75mg/L ox-LDL for 24h, followed by treatment with Euonymine (60, 90,120μM) for 24 h. Thereafter, VSMCs were subjected to staining with Giemsa. Early apoptosis and cell necrosis of VSMCs induced by subjection to different concentrations of Euonymine. Nuclear hyperchromat in staining, fragmentation, chromatin accumulation, and mass distribution could be observed around the nucleus, a small number of cells could be observed to be swollen, and apoptotic bodies surrounded by complete membranes could be observed around VSMCs, compared to the negative control group. The red arrow indicates marked morphological changes of apoptosis. B, Morphological presentation of apoptosis of VSMCs induced by Euonymine treatment (TUNEL staining, ×100 magnification). a, Representative image depicting morphology of apoptosis in VSMCs induced by Euonymine treatment. b, Quantitative data have been presented as the percentage of TUNEL-positive cells derived from the total number of cells. Cells were harvested 72 h after the addition of different concentrations of Euonymine. Data have been presented as mean ± SD (n = 6). *P <0.05 vs. control, ^#^P <0.05 vs. ox-LDL. C, Transmission electron microscopy (TEM) was performed. Black arrows indicate ultrastructural changes of VSMCs induced by Euonymine treatment. a. Normal VSMCs with affluent chromatin and marked nucleus(×4000 magnification). b. Earlier apoptotic VSMCs with chromatin condensation, chromatin pyknosis, fracture (×5000 magnification). c. Crescent-shaped chromatin merge together under the membrane of the nucleus (×5000 magnification) d. Apoptotic VSMCs with an apoptotic body (×5000 magnification). D and E, Flow cytometry analysis performed to detect the effect of apoptosis in VSMCs.

### 2.8 Euonymine promotes apoptosis of VSMCs via the mTOR pathway *in vitro*

This investigation was conducted to further explore the molecular mechanism of Euonymine-mediated apoptosis induction in VSMCs. We determined the expression of downstream-related apoptotic factors of PTEN/AKT1, including mTOR, P-mTOR, Bax, Bcl-2, cleaved PARP, cytochrome C. We found that exposure of VSMCs to Euonymine resulted in a dose-dependent increase in the cleavage of caspase-3, Bax/Bcl-2, as shown in Figure10 A. Bax/Bcl-2 ratio was evaluated to reflect the apoptosis level, and the results indicated that cell apoptosis was inhibited by ox-LDL. However, cell apoptosis was induced by Euonymine treatment. Notably, ox-LDL inhibited the release of cytochrome C from the mitochondria to the cytoplasm, whereas Euonymine treatment reversed this process, as shown in Figure10. B. Cleaved poly (ADP-ribose) polymerase (PARP), a substrate for caspase-3, is a well-known marker of apoptosis and its level increases with the concentration of Euonymine, as shown in Figure10.C. Based on the above-mentioned results, it is suggested that Euonymine inhibits proliferation, migration, and promotes apoptosis of VSMCs through a mitochondria-dependent pathway.

**Figure 10.**
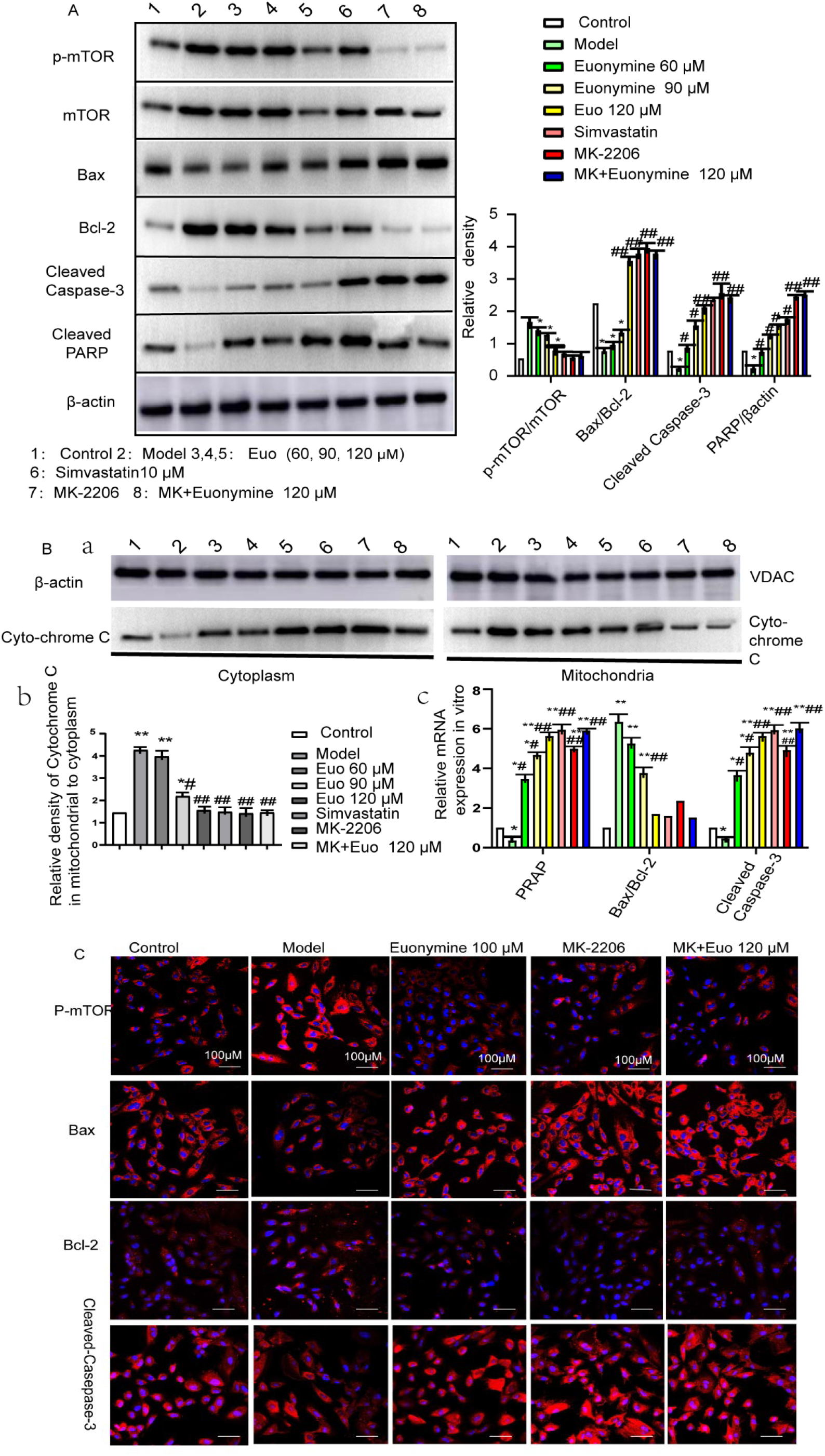
Euonymine triggered apoptosis of VSMCs *in vitro*. Serum-starved VSMCs were pre-treated with Euonymine (60, 90, 120 μM) for 24 followed by treatment with 75 mg/L ox-LDL for 24h. In addition, VSMCs were treated with MK-2206 30μM in the absence or presence of Euonymine (120μM) for 24h. A and B, Expression levels of indicated proteins and genes were evaluated by western blot assay and RT-PCR. C, Immunofluorescence images of indicated proteins. The data are expressed as the means ± SDs. *P< 0.05, and ** P< 0.01 vs. the control group; ^#^P < 0.05 and ^##^P < 0.01 vs. the ox-LDL group. n=6. β-actin was used as the internal reference, and VDAC as mitochondria markaer. Scale bar=100μm.

### 2.9 Euonymine induces G0/G1 phase cell cycle arrest in a p38/MAPK dependent manner

To further elucidate the mechanism of Euonymine-induced apoptosis in VSMCs, the cellular DNA distribution was detected via propidium iodide (PI) staining. Our results showed that compared to the control group, the number of cells in the G0/G1 phase increased in the Euonymine group, while the expression of cell cycle proteins CyclinE and CDK2 was significantly lower in the Euonymine group compared to that observed in the model group. Our experimental results confirmed that mitotic blockade in the G0/G1 phase in VSMCs of the Euonymine group was closely related to the expression of cyclinE1 and CDK2 of cell cycle proteins, as shown in Figure 11. A and B.

**Figure 11.**
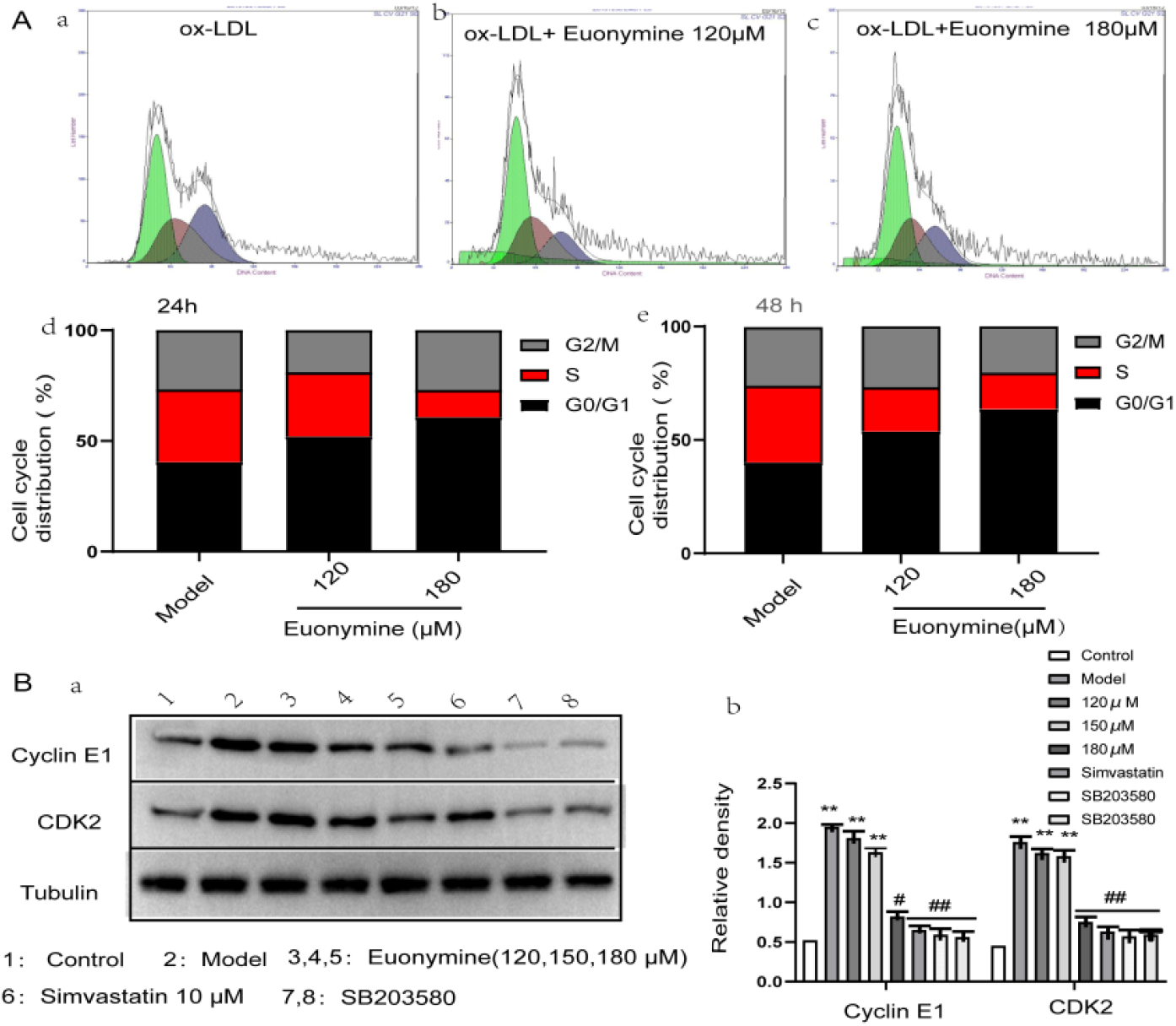
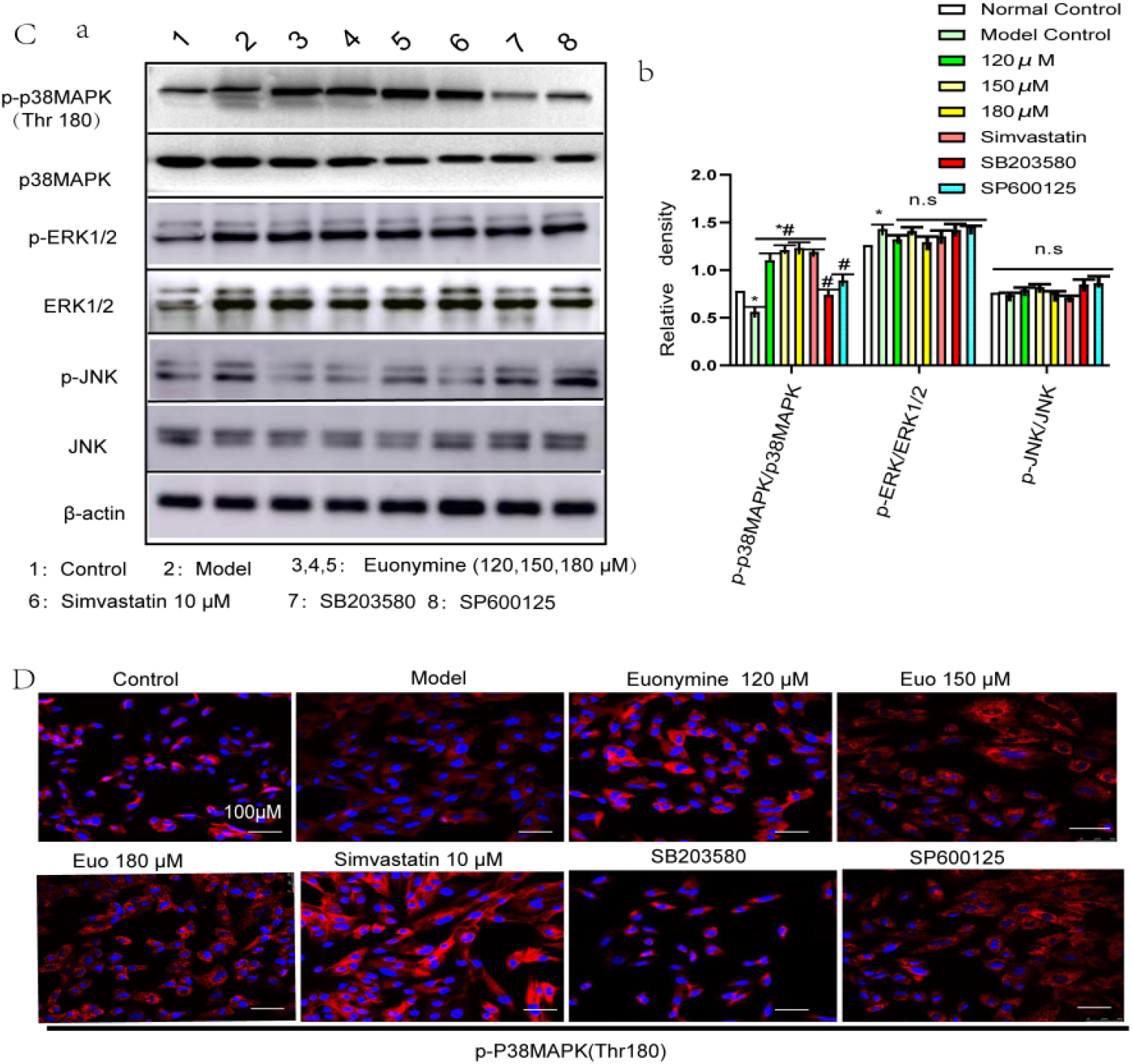
Euonymine induces G0/G1 phase cell cycle arrest in a p38/MAPK dependent manner. VSMCS were pre-treated with 10µM SB203580 (a p38MAPK specific inhibitor) or SP600125 (a JNK specific inhibitor) for 1 h followed by treatment with Euonymine (120-180µM) and ox-LDL(75mg/L) 24h, respectively. A, Representative measurement of DNA distribution in serum-stimulated VSMCs in absence or presence of Euonymine. (a,b,c), DNA histogram of VSMC treated with different concentrations of Euonymine for 24 hours. (d,e), Cell cycle distribution of VSMCs treated with Euonymine for 24 h and 48 h. B and C, Expression levels of indicated proteins were evaluated by western blot assay. D, Immunofluorescence images of indicated p-p38MAPK(Thr308). The data are expressed as the means ± SDs. *P< 0.05, and ** P< 0.01 vs. the control group; ^#^P < 0.05 and ^##^P < 0.01 vs. the ox-LDL group. n=6. Tubulin was used as a loading control; n.s. represents no significant difference.

Considering that Euonymine inhibits proliferation and migration, as well as induces a G0/G1 phase blockade and apoptosis in VSMCs, previous studies have shown that the P38MAPK signaling pathway plays a key role in cell cycle progression, along with cell invasion and migration (27). Thereafter, the expression levels of p38/MAPK pathway-related proteins (Thr180) were detected via WB and IF. The results demonstrated that the phosphorylation level of p38/MAPK was significantly decreased after 24 h incubation of ox-LDL with VSMCs, as shown in Figure11C. p38MAPK/JNK phosphorylation level was significantly increased in a dose-dependent manner after Euonymine treatment, while the phosphorylation status of JNK and ERK1/2 did not change significantly. To confirm the role of p38/MAPK and JNK/MAPK pathways in the inhibitory effect of Euonymine on VSMCs, we used the p38MAPK-specific inhibitor SB203580 and the JNK/MAPK-specific inhibitor SP600125. Western blotting analysis showed that the SB203580 group presented with reversal of the elevated expression of p38 phosphorylation in ox-LDL-treated cells, while the expression of JNK and ERK1/2 in the SP600125 group was not significantly different. These results suggested that p38/MAPK might be involved in the anti-proliferation effect of Euonymine, but it might not play roles in the JNK/MAPK pathway. Subsequently, we investigated the correlation between p38/MAPK and induced G0/G1 phase blockade. The expression of cyclin E1 and CDK2 was found to be significantly lower after investigating VSMCs without treatment and with p38MAPK-specific inhibitor SB203580 treatment, compared to the model group; additionally, reversal of the high expression of cyclin E1 and CDK2 was induced in the presence of ox-LDL. Taken together, our data demonstrated that Euonymine induced G0/G1 phase cell cycle arrest in a p38/MAPK-dependent manner and not in a JNK/MAPK-dependent manner.

## 3. Discussion

Patients with atherosclerosis are at higher risk of developing restenosis in the vascular bed, and human PCI procedures are often performed in patients with comorbidities such as atherosclerosis. However, ISR can occur due to the progressive thickening of the neointimal hyperplasia within the stent into an unstable AS plaque after PCI, which may rupture and lead to the occurrence of an acute adverse cardiovascular event, considerable affecting patient prognosis[33]. Therefore, we used the rabbit carotid balloon injury and porcine atherosclerotic coronary implantation models to simulate the dynamic pathological progression of ISR after PCI in humans. We used the rabbit carotid balloon injury model, wherein the main pathological process simulates intimal hyperplasia development after infliction of balloon dilation injury, while the pathological process of the porcine atherosclerotic coronary implantation model encompasses the pathological process of mechanical injury inflicted by stent dilation and intimal hyperplasia development after the occurrence of stent injury; the pathological changes of ISR caused by intimal hyperplasia development in the two animal models are similar to the clinical/pathological process of ISR after PCI. The dynamic development process of ISR in the two animal models is similar to the clinical pathology of ISR after PCI, which presents with avaluable clinical reference value. In a rabbit carotid balloon injury model, HE staining was performed and revealed that Euonymine significantly reduced the extent of neointimal hyperplasia after infliction of arterial injury. Meanwhile, the results of WB and IF suggested that angioplasty caused upregulation of expression of synthetic markers (osteopontin) and downregulation of expression of contractile markers (SM22α, αSMA), which were attenuated by Euonymine treatment, as shown in Figure 5. A and B. In summary, our data indicated the effectiveness of Euonymine treatmentin balloon injury-induced neointimal hyperplasia. Additionally, we evaluated the effect of Euonymine treatment on post-PCI ISR by implementing HE, QCA, and OCT techniques in a preclinical porcine coronary artery stent model. The results of the three experiments corroborated each other and highlighted the extent of lesions in vascular neointimal hyperplasia. Additionally, the findings helped visually confirm that coronary intimal hyperplasia was effectively suppressed in the ISR stent model after implantation of Euonymine-coated stents and showed good histocompatibility and safety, as shown in Figures 2-4. OCT is an *in vivo* test that can help compensate for QCA and has garnered attention as the new gold standard for intravascular imaging of stents, atherosclerotic progression, vulnerable plaques, and neointimal hyperplasia(26). The above-mentioned experimental results suggest that Euonymine significantly reduces the extent of neointimal hyperplasia after arterial injury, thus providing robust evidence for the development of a novel stent-coated drug for clinical application in ISR.

To further explore the mechanism of Euonymine-mediated inhibition of neointimal hyperplasia, we performed *in vitro* experiments. ISR occurrence after PCI is a complex pathophysiological process involving multiple factors and components (28). Recent studies have shown that excessive proliferation, migration, and insufficient apoptosis of VSMCs constitute the main causes of neointima hyperplasia development(29–31). Under normal physiological conditions, VSMCs constitute a highly specialized and almost quiescent cell population whose main function is to maintain vascular tone and to ensure vasoconstrictor function. PCI leads to neointimal damage and the exposure of the damaged intimal material to blood, resulting in platelet activation, release of various active factors, and changes in intercellular composition, thereby mediating excessive proliferation of VSMCs and migration from the media to the intima (32–34). Additionally, VSMCs undergo phenotypic changes and differentiate from cells with contractile function to cells with secretory function, which secrete extracellular matrix and form the neointima, covering the inner and outer surfaces of the stent and stent wall, resulting in the thickening of the neointima of the vessel; they may invade the lumen of the vessel and cause a narrowing of the lumen(35), a phenomenon which is consistent with that reported in our study. After Euonymine intervention, we found that ox-LDL caused up regulation of expression of synthetic markers (osteopontin) and down regulation of expression of contractile markers (SM22α, αSMA), conditions which were found to be reversed subsequently by Euonymine treatment *in vitro* models, as shown in Figure 6. A and B. Concomitantly, our findings demonstrated that Euonymine inhibited the proliferation and migration of VSMCs *in vitro*, as shown in Figure 7. We tentatively suggest that Euonymine inhibits neointimal hyperplasia partly by inducing the formation of a contractile phenotype of vascular smooth muscle cells and by inhibiting the proliferation and migration of VSMCs, thereby promoting rapid vascular repair.

To further ascertain the effect of Euonymine on ISR, we investigated its specific molecular mechanisms. First, we evaluated the effects of Euonymine on PTEN/AKT/mTOR and apoptotic factors in the injured arterial wall of rabbits, focusing specifically on changes in the expression of the phosphorylation site AKT1 (Ser437), to provide evidence of a mechanism underlying the inhibition of VSMC proliferation, migration, and induction of apoptosis. It has been demonstrated that AKT1 is involved in the proliferation of neointimal hyperplasia of pulmonary arteries through smooth muscle cell activation, which promotes the proliferation and migration of VSMCs (36, 37). Additionally, through the siRNA of AKT1, Diosgenin inhibits cell proliferation and induces apoptosis, thus demonstrating activities as an anticancer agent (38). Our findings were consistent with those reported in previous studies, indicating that the level of P-PAKT1 (Ser437) phosphorylation in rabbit common carotid arteries was significantly increased on day 14 after balloon injury compared to uninjured arteries and that Euonymine treatment reversed the progression of this pathological process, as shown in Figure 6 C and D. The results of WB, HE, and IHC experiments indicate that Euonymine local perfusion reduces the extent of neointimal hyperplasia, as shown in Figure 2 and Figure 6. In the present study, we also examined the level of Euonymine-mediated inhibition of AKT1 phosphorylation *in vitro* and VSMCs were subjected to pre-treatment with an AKT1 inhibitor MK-2206. Our study showed that Euonymine treatment inhibited VSMC phenotypic transformation by targeting AKT1 expression as shown in Figure 7, a process that was mechanistically mediated through the typical PTEN/AKT1/mTOR pathway and its downstream apoptotic signaling pathway, indicating consistency with the previously reported findings(18). In conclusion, Euonymine inhibits the phenotypic transition of VSMCs by targeting AKT1 to mediate the proliferation and migration of VSMCs, thereby inhibiting neointimal hyperplasia development.

Mitogen-activated protein kinase (MAPK) exhibits responses to extracellular stimuli and participates in intracellular signaling, thus playing an important role in the regulation of cell proliferation, differentiation, and apoptosis(39). p38/MAPK has been revealed to play a key role in the DNA damage response and, once activated, may lead to cell growth inhibition and apoptosis (40). The p38/MAPK signaling pathway has been confirmed to play a key role in cell invasion and migration (41). Our study found that Euonymine treatment might promote apoptosis in VSMCs partially by enhancing the p38MAPK-associated mitochondria-dependent apoptotic pathway, as shown in Figures Figure10. C and D. However, subsequent reports suggest that JNK/MAPK also demon stratesan anti-apoptotic function that supports cell survival and growth(42). These complex functions regulated by MAPKs may be determined by investigating the stimuli involved, cell type, and duration of activation. Based on the information presented in previous and latest studies, our findings provide crucial evidence that p38MAPK may be a potent target for the treatment of ISR. Thus, the combined in vivo and in vitro data suggest that Euonymine inhibits phenotypic transition of VSMCs and suppresses the activation of AKT1 phosphorylation sites, and activates the P38AMPK pathway, but exerts no significant effect on JNK and ERK1/2 expression in VSMCs. Additionally, we found that Euonymine demonstrated more marked functions as a potent inhibitor of AKT1 expression rather than an activator of P38MAPK. For example, treatment with 120 μM Euonymine inhibited AKT1 expression (Figures 7 and 8); however, a concentration of 180 μM was necessary for the activation of p38MAPK, as shown in Figure11. Therefore, we suggested that inhibition of AKT1 phosphorylation site activation might be the main mechanism by which Euonymine inhibited VSMC proliferation and migration and attenuated intimal hyperplasia proliferation.

Presently, in terms of the mechanism of ISR, excessive proliferation and migration of smooth muscle cells, which leads to a cell proliferation-apoptosis imbalance and subsequent neointimal hyperplasia development, have been largely considered as the main cause of ISR(43, 44). To further explore the mechanism of anti-neointimal hyperplasia of Euonymine, we explored Euonymine-induced apoptosis of VSMCs in a holistic animal model and a cell injury model. Both *in vitro* and *in vivo* data showed balloon injury and ox-LDL-induced down regulation of Bax/Bcl-2 ratio and inactivation of caspase 3 and PARP. It has been shown that the pro-apoptotic factor Bax promotes the release of cytochrome C and activates caspase-3 to execute the apoptotic program, whereas the anti-apoptotic molecule Bcl-2 inhibits the activation of Bax and release of cytochrome C to suppress apoptosis (43). Additionally, our results were consistent with those reported in previous studies. Our study showed that the events of elevation of mitochondrial cytochrome C levels and decrease in the levels of cytoplasmic cytochrome C were both reversed after Euonymine treatment, as shown in Figure 10. Therefore, we suggested that Euonymine induced apoptosis in VSMCs via the p38MAPK-related mitochondria-dependent apoptotic pathway, which might be considered an additional explanation for the attenuation of neointimal hyperplasia.

In the present study, using *in vitro* and *in vivo* models, we demonstrated that Euonymine drug-eluting stents inhibited in-stent restenosis by targeting AKT1 and p38MAPK to enhance the contractile phenotype of VSMCs to prevent neointimal hyperplasia development. Therefore, Euonymine may be considered a potential molecule for application in the prevention and treatment of ISR after PCI.

Notably, the study presented with certain limitations. The effects of Euonymine in carotid arteries in animal models may differ from those exerted in diseased human vessels, which may limit the extrapolation of the results of the present study to human outcomes. It should also be noted that our ability to translate effective concentrations *in vitro* to effective concentrations *in vivo* was limited. Nevertheless, we observed a significant inhibition of endothelial proliferation via Euonymine treatment in animal models of injured arteries, suggesting that the concentrations used *in vivo* generalize the *in vitro* effects of Euonymine. While previous studies have demonstrated that delayed endothelial healing is closely associated with late thrombosis in drug-eluting stents, the Euonymine-coated stent-implanted segment coronary arteries presented with intact endothelial cell coverage and did not exhibit stent thrombosis at 28 days post-procedure, suggesting good histocompatibility and safety of Euonymine-coated stents despite the short period observed. Furthermore, the determination of inhibitory effects of Euonymine on vascular neointima through PTEN/AKT/mTOR and P38MAPK, and the exploration of a possible existence of an association between the two, will be the future aims of our study.

## 4. Methods

### 4.1 Chemicals and reagents

Due to word limits, this section is presented as supplementary material.

### 4.2 Preparation of Euonymine solutions

Euonymine was purchased from the Kunming Institute of Botany, Chinese Academy of Sciences, and with a purity of 99.5% (Kunming, China).

#### 4.2.1 Euonymine preparation for whole animal experiments

Considering a concentration of0.5mg/mL as an example, 5mg Euonymine was dissolved in 25μL DMSO and the volume was adjusted to 10mL, such that the final concentration of Euonymine was 0.5mg/mL (the final concentration of DMSO was 0.25%). The solution of the above-mentioned concentration of 0.5 mg/mL (the 0.5 mg/mL dose group) was diluted to obtain0.25 mg/mL Euonymine solution.

#### 4.2.2 Euonymine preparation for cell experiments

Euonymine (80.5783g) was dissolved in 2.9 mL of dimethylsulfoxide (DMSO) to obtain a stock solution of Euonymine with aconcentration of 1 × 10^5^ μM, and it was stored in the dark at 4°C. The stock solution was diluted to the required concentrations (7.5, 15, 30, 60, 90, 120…μM) using the Dulbecco’s modified Eagle medium (DMEM).

#### 4.2.3 Euonymine drug-eluting stent development

The following stents were developed and provided by Shandong Ruiantai Medical Technology Co., Ltd.; the Bare-metal stent (BMS) group, three Euonymine-eluting stent groups (0.14 [low-dose group], 0.70 [medium-dose group], and 2.80 [high-dose group] μg/mm^2^), and the Sirolimus-eluting stent (SES) group (1.40μg/mm^2^) were established. All stents were composed of 316L stainless steel, and the surface coating was composed of coated drugs and degradable poly lactic-glycolic acid (PLGA). The unified specification of the stent is 3.0mm×18mm (diameter × length). The stent was sterilized by ethylene oxide treatment and was properly packaged. The stent was designed with an open-loop structure, cut from 316L stainless steel material, with a stent beam thickness of 0.0035 inches.

### 4.3 Animals and the establishment of animal models

#### 4.3.1 Animals

Twenty New Zealand rabbits of either gender, weighing 2.0-2.5 kg, and 20 male healthy minipigs were purchased from the Animal Center of Kunming Medical University. The production license numberis SCXK (Dian)K 2015-0002, and the experimental animal license number is SYXK(Dian) 2017-0006. The animals were exposed to a 12-h light/dark cycle, and they had free access to food and water. All procedures involving minipigs were approved by the Animal Research Ethics Committee of Kunming Medical University (animal ethics license number: KMMU2018219).

#### 4.3.2 Rabbit carotid artery balloon angioplasty modeling, drug administration, sampling, and pathological examination

##### 4.3.2.1 Rabbit carotid artery balloon angioplasty modeling, drug administration, sampling

According to the information present in the existing literature (45, 46), the white rabbits were subjected to anesthetizationusing 5% sodium pentobarbital (0.5 mL/kg body weight) via ear margin intravenous administration until the disappearance of the eyelash reflex. The neck was routinely debrided and disinfected, following which the skin was incised 3-4 cm in the direction of the trachea in the middle of the neck, and the subcutaneous fascia and muscles were separated successively; the common carotid artery and the bifurcation of the external and internal carotid arteries were investigated for medial to the left sternocleidomastoid muscle, and the proximal end of the common carotid artery was temporarily blocked using an arterial clip; additionally, sodium heparin was administered intravenously at a dose of 200 U/Kg body weight and then the distal end of the external carotid artery and the peripheral common carotid artery were ligated. A small incision was introduced at the distal end of the external carotid artery using ophthalmic scissors, and a 0.014-inch soft-tipped guidewire was placed into the superior cavernous artery under the guidance of a guiding needle, following which a 2.5×10 mm^2^ diameter balloon catheter was introduced in a retrograde manner along the guidewire to the proximal end of the common carotid artery (approximately 4-5 cm in length); the balloon was filled with saline (2 atm) and was slowly pulled back against resistance to the external carotid artery incision, following which it was evacuated. The catheter was then returned, and the procedure was repeated once in each of the three directions 120° apart to fully debride the common carotid artery intima. The balloon and guidewire were withdrawn, the external carotid artery was ligated, and 200,000 units of penicillin were administered locally. In the sham-operated group, only the external carotid artery was ligated and no balloon was inserted.

##### 4.3.2.2 Drug administration

The rabbits were randomly divided into the following four groups: (1) sham-operated group. Ligation of the left external carotid artery was performed only; (2) the model group. After left carotid artery balloon angioplasty was performed, local intraluminal infusion of normal saline containing 0.25% DMSO was conducted for 30 min. (3) and (4) the Euonymine group with local delivery of the drug(0.25 mg/mL, 0.5mg/mL): immediately after operation, the local lumen was perfused with 0.25 mg/mL or 0.5 mg/mL Euonymine for 30 min.

##### 4.3.2.3 Local infusion of Euonymine

After the intraoperative balloon catheter was used to completely strip the endothelium of the common carotid artery, the balloon was withdrawn at this juncture, the guiding needle was carefully introduced along the guiding wire to the distal end of the common carotid artery, and the external carotid artery incision was ligated, following which the proximal end of the internal and common carotid arteries were clamped using an arterial clip concurrently; the blood in the vascular lumen was with drawn using a 2 mL syringe from the end of the guiding needle, following which saline was introduced and the area was flushed twice. Then, 0.25 mg/mL or 0.5mg/mL of the Euonymine solution (500μL) was introduced. After 30 min, the solution was withdrawn, saline was introduced and the area was flushed twice, and the guide needle and arterial clip were withdrawn to restore blood flow and one time per day. Animals were sacrificed on days 14 and 28 post angioplasty, and skin and subcutaneous tissues of the neck were excised, following which the left common carotid artery was isolated and the injured segment of the common carotid artery was intercepted for 4 cm. After washing off the residual blood in the lumen with saline in a Petri dish, the segment was directly fixed in a 4% paraformaldehyde phosphate buffer solution.

##### 4.3.2.4 Pathological examination (hematoxylin and eosin staining and immunohistochemical assay)

Hematoxylin and eosin (HE) staining: The vessels were subjected to fixationin 4% paraformaldehyde phosphate buffer solution. After rinsing with tap water, a section of the fixed tissue was routinely dehydrated to obtain a transparent specimen, following which it was impregnated with wax, embedded in paraffin, and sectioned intermittently. Three sections of each specimen were acquired at distal, middle, and proximal sites. Paraffin-embedded sections were selected from days 14 and 28 and were stained with HE, followed by subjection to LEICAQ550CW Image analysis.

Immunohistochemical (IHC) assay: Paraffin-embedded sections were stained 14days postoperatively, following which slices were dewaxed using water and were incubated with 3% hydrogen peroxide-methanol solution for 10 min at room temperature; antigen was thermally retrieved. The sections were incubated with 5% to 10% normal goat serum; primary antibody (mouse anti-PCNA monoclonal antibody, 1:50 v/v) was added in a dropwise manner. Following overnight incubation at 4°C and subjection to washing steps with phosphate buffer (PBS), biotin-labeled secondary antibody solution was added in a dropwise manner and incubation was performed for 1 h at 37°C in a wet box; subsequently, rinsing steps were performed with PBS. Horseradish enzyme-labeled streptavidin was added and incubation was conducted at 37°C for 1 h in a wet box, followed by subjection to washing steps using PBS. Thereafter, DAB color development, rinsing steps with tap water, re-staining experiments, dehydration, achievement of translucency, and sealing steps were observed. Each section was randomly selected under a high magnification of alight microscope. Six fields of view were randomly selected for each section under high magnification of a light microscope, and the total number of intima and media VSMCs were counted separately. The average percentage was calculated as the PCNA-positive cell index (PI).

#### 4.3.3 Establishment of porcine coronary artery stent model, sample collection, and pathological examination

##### 4.3.3.1 Establishment of porcine coronary artery stent model

I. Animal grouping: Twenty minipigs were randomly divided into Bare metal stent (BMS), three Euonymine-eluting Stent groups (0.14 [low-dose], 0.70 [medium-dose], and 2.80 [high-dose] μg/mm^2^), and Sirolimus-eluting Stent (0.14μg/mm^2^) group. A model of porcine atherosclerosis was replicated according to the previous research of our research team[25], and then a stent was placed in the coronary stenosis site, and left or right coronary angiography and OCT were performed through the right femoral artery or left femoral artery route after stent implantation and 28 days, (See later for details of the specific operation).

II. Pre-operative preparation. III. Femoral artery puncture or femoral artery dissection. IV. Coronary angiography and stent implantation. V. Surgical puncture site handling. VI. Postoperative treatment VII. Coronary angiography and analysis. VIII. OCT examination and analysis. Due to word limits, this modeling section is presented as supplementary material.

##### 4.3.3.2 Collection of pathological samples and HE staining

The minipigs were sacrificed 28 days after the operation, and the blood vessel specimens of the stent segment were collected. After bleeding and executing the animal after the completion of imaging, the heart was removed by opening the chest cavity, preserving the ascending aorta, and was immersed in saline. A dilatation tube was inserted from the ascending aorta to the left and right coronary artery openings, and the coronary artery was repeatedly flushed with saline 5-6 times, following which 10% formaldehyde solution was perfused from the aortic root at a pressure of 70 mmHg for 30 min. The stent implanted segment was carefully separated and placed in a 10% neutral formaldehyde fixative for the preparation of light microscopy specimens.

The stented coronary artery specimens were soaked in 10% neutral formaldehyde solution for more than 24 h. A fixed section of the vessel was selected, following which it was rinsed with tap water, dehydrated, embedded, sectioned, and degreased; then, interrupted serial sections were stained with hematoxylin-eosin to observe the pathological changes of vessel histology under the microscope; three cases were randomly observed in each group, and five high magnification fields were randomly selected for each section.

##### 4.3.3.3 Evaluation of morphological parameters of restenosis in the stented segment

The neointimal thickness, residual lumen area, internal elastic plate surrounding area, neointimal area, and percentage of intimal stenosis area were determined and calculated by using the HMIAS-2000 image analysis software.

### 4.4 Cell isolation and culture

Rabbit thoracic aorta primary VSMCs were isolated from the thoracic aorta by adopting an enzymatic digestion technique and the cells were cultured in DMEM according to the protocols reported in the literature (47, 48). Briefly, the thoracic aorta was isolated, adventitial tissue was removed, DMEM culture medium containing 20% fetal bovine serum was added, and incubation was performed at 37°C and 5% CO_2_ in an incubator for 2 h, with a solution containing penicillin-streptomycin (100 U/mL penicillin and 100 μg/mL streptomycin). Then, cells were passaged, a small amount of sterile 0.1% PBS solution was added to subject the cells to washing steps, following which elastase (1 U/mL, LS006365) digestion solution was added to perform digestion of the cells; blown repeatedly to disperse the cells into a single cell. Subsequently, the single-cell suspension was divided into new culture flasks and was replenished with the culture medium. The morphological observation via phase-contrast microscopy and anti-α-actin monoclonal antibody staining confirmed the presence of VSMCs. The cells were stored under freezing conditions, and cells belonging to the 3rd-5th generation were used for experiments.

### 4.5 Cell viability and proliferation assay

The cell growth curve was determined by performing MTT assay. The VSMCs were used in the experiment. The 3-5th generation VSMCs were used in the experiment. The cells were subjected to treatment with DMEM 12 h before the assay. After subjection to various treatments, 20 μL of MTT was added to each well and the plates were incubated for 2.5 h. Thereafter, we measured the absorbance of the samples using Multiskan (Thermo) at 570 nm. The relative expansion rate was computed by considering the absorbance proportion of the drug-treated group to the control group.

### 4.6 Euonymine treatment *in vitro*

The experimental groups established were as follows: normal control group; ox-LDL (75 mg/L) injury model group;③simvastatin-positive control group (final concentration of 10μmol/L); ox-LDL+different concentrations of Euonymine (7.5,15,30,60,90,120…μM); ox-LDL+Euonymine (120 µM) and MK-2206 (an inhibitor of the AKT1phosphorylation site); and ox-LDL+MK-2206. The cell suspension was prepared by selecting the 6th-8th generation of VSMCs with good growth characteristics and was cultured in the CO2 incubator at 37℃, 5% CO2, and 95% saturation humidity for 24h; then, the original culture medium was discarded and the same volume of serum-free medium was added for assimilation for 24 h. In the normal control group, the complete medium was added, and in the other groups, 75 mg/L ox-LDL medium was added to continue the culture. After 24 h, the cells were collected and the culture medium was removed via washing. Total protein or RNA extraction was conducted after cell lysis was performed by adding the lysis buffer, follow-up experiments were conducted. There were 6 parallel holes in each group, and the experiment was repeated 3 times.

### 4.7 Wound healing assay

Cell migration was detected by conducting an *in vitro* wound healing test, and experiments were grouped as above. VSMCs were added in Culture Insert-2 Well (Ibidi, No: 81176, Martinsried, Germany) at a density of 5×10^4^ cells/mL with 80 uL cell suspension per well. After incubation in an incubator with 95% saturation humidity for 24 h, each group was incubated with 75 mg/L of ox-LDL. The normal group was incubated with an equal volume of the medium and was subjected to serum starvation for 24 h. The original culture medium was discarded, following which a fresh complete culture medium was added to the normal and model groups, culture medium with serial concentrations of Euonymine was added to the experimental group, and culture medium containing 10µM simvastatin was added to the positive control group; incubation was continued in the incubator for 24 h. When the cells reached 100% confluency, we removed the inserts to create a cell-free interstitial space at an equal distance of 500 μm (please refer to the manufacturer’s instructions for specific experimental steps). Cells were subjected to washing steps with PBS to remove non-adherent cells. Additionally, we imaged live cells (0 h) and acquired photographs (×100 magnification). Then, to each group, the complete culture medium (serum < 2%) was added and incubation was continued for 24 h. PBS was removed by conducting washing steps twice, and the results were photographed and recorded as findings obtained at 24h. Cell migration distance (d) was expressed as 0-h distance subtracted from 24-h distance in um, and the experiment was repeated thrice.

### 4.8 Wright-Giemsa staining

The VSMCs suspension was centrifuged and resus pended in PBS to obtain a cell concentration ranging from 10^5^-10^6^/mL. The cell suspension was collected for flaking, following which it was subjected to drying conditions and was fixed in methanol for 5 min. Giemsa staining solutions were added to the cells, and staining was performed at room temperature for 10-20 min (to observe the degree of staining under the microscope); the slides were immersed in xylene for 3 min and were sealed with resin after the achievement of transparency. The nuclear morphology of the cells was observed under a light microscope, and photographic and image scanning analyses were performed. Five images of each group of cells were randomly selected, and the apoptosis rate of each group was calculated.

### 4.9 TUNEL staining for DNA fragmentation analysis

In situ detection of DNA lysis fragments was performed by implementing the TUNEL method. According to the above-mentioned grouping and drug administration method, the cells of each group were collected after 72h of incubation and cell suspensions were prepared, followed by subjection to shaking conditions. The experimental procedures were performed according to the kit instructions as follows: (1) 40g/L paraformaldehyde was added for fixationat room temperature for 30min; (2) 0.3% (v/v) hydrogen peroxide methanol solution was added for 30min to block the endogenous enzyme activity; (3) the TUNEL reaction mixture was added and incubation was performed at 37°C for 60min; (4) the TUNEL reaction mixture was added and incubation was performed for 60min; transformant-POD was added and incubation was performed at 37°C for 30min; (5)the DAB substrate was added and incubation was performed at 37°C for 10 min. The apoptotic cells exhibited the presence of brownish-yellow granules in the nucleus, along with chromatin aggregation, nucleus consolidation, and fragmentation into several granules; the negative control did not show the presence of colored granules, and apoptosis index (AI) was calculated. Quantitative data have been expressed as the percentage of TUNEL-positive cells derived from the total number of cells.

### 4.10 Ultrastructural observation via transmission electron microscopy

VSMCs were subjected to treatment with different concentrations of Euonymine for 72h, and then 1×10^6^ cells from each group were subjected to centrifugation and precipitation. After supernatant removal, specimens with the cells were routinely fixed, dehydrated, permeabilized, embedded, sectioned to obtain an ultra-thin sample, and were double stained with uranium-lead, following which observation and photography under transmission electron microscopy were conducted.

### 4.11 TUNEL/PI assay

Serum-starved VSMCs were incubated with 75mg/L ox-LDL for 24h, followed by treatment with Euonymine (60, 120μM) for 24 h. Cells were collected and incubated in 1mL cold PBS to prepare a cell suspension with 1 × 10^6^ cells, followed by centrifugation at 4°C for 10 min at a speed of 1000r/min; the supernatant was discarded, and cells were resuspended in 200µL Rinse Buffer. Then,10µL of FITC-the dUTP labeling mixture and 5µL of PI were used and mixed gently, and the mixture was subjected to reaction for 15 min at room temperature with closed light or 30 min at 4°C with protected light; 300µL of Rinse Buffer and samples were added to the spiking chamber of FCM within 1 h, and the green fluorescence of FITC-dUTP was measured at 520±20nm and the red fluorescence of PI was measured at > 620nm; 20,000 cells per sample were counted, following which the data were illustrated using graphs generated automatically via a computer and the apoptosis rate was determined by using the Cell Quest software.

### 4.12 Flow cytometry analysis

Prepared 1×10^6^ VSMCs cell suspensions were centrifuged at 1000r/min for 5 min, followed by subjection to washing steps with PBS. The cell suspensions were subjected to overnight fixation at 4℃ with 70% ethanol pre-chilled at −20℃, followed by subjection to washing steps twice with PBS at 4℃; RNase A was added to a final concentration of 50 mg/L, and incubation was performed in a water bath at 37℃ for 30 min. The final concentration of 50 mg/L PI was used for incubation at 4℃ for 30 min. The fluorescence intensity was measured via flow cytometry, and the DNA content was analyzed via flow cytometry.

### 4.13 Western blotting (WB)

Western blotting was implemented to quantitate protein levels of PI3K, P-PI3K, AKT1, P-AKT, Bcl-2, Bax, mTOR, p-mTOR, cleaved Caspase-3, α-SMA, SM22α, and osteopontin in carotid tissue and ox-LDL-induced cell model. Protein concentrations of tissues and cells were determined by using a protein assay kit. Samples of supernatants containing 40µg protein of tissues and cells were loaded and were heated to 95℃ for 10 min, following which they were subjected to separation via sodium dodecyl sulfate-polyacrylamide gel electrophoresis using 8% or 10% gels and the Mini-Protein II apparatus (Bio-Rad, CA, United States). Protein bands were electroblotted onto the polyvinylidene difluoride (PVDF) membrane. After subjection to blockade in a 5% BSA solution in Tris-buffered saline with Tween 20(TBST) for 1.5h, the membranes were incubated with [PI3K, P-PI3K, AKT(1:2000), P-AKT1(Ser437), P-AKT1(Thr308), SM22α and α-SMA, OPN, Bcl-2, Bax(1:1500), mTOR, p-mTOR(1:2000), cleaved Caspase-3, (1:2500), β-actin (1:1000), GAPDH(1:1200), primary antibodies overnight at 4℃. After subjection to rinsing steps thrice with TBST, membranes were incubated with secondary antibodies, either with horseradish peroxidase-conjugated anti-rabbit IgG (1:2000), for 1h prior to subjection to washing steps and development with ECL reagents. The chemiluminescence signal was imaged using the ChemiDoc XRS system (Bio-Rad), and protein band signals were quantified by using the ImageJ 1.49v Software. The experiment was repeated three times.

### 4.14 Real-time PCR (RT-PCR) analysis

To investigate the effects of Euonymine on the vascular tissue of stent segment and cellular models, RT-PCR was also performed to quantitate mRNA levels of Bcl-2, Bax, Caspase-3, and mTOR. The TRIazol reagent kit was used to perform total RNA extraction from vascular tissue and cells, and reverse transcription was performed using the Quant Script RT kit. PCR was performed using ABI7900HT (ABI Company, USA) according to established methods. 2^−ΔΔCT^ values were used to indicate the quantity of relative expression of five types of RNA (mTOR, Bcl-2, Bax, and cleaved Caspase-3). Five PCR primers were selected randomly and prepared to analyze the changes in carotid tissue and cells. The PCR conditions were as follows: thermal denaturation 95°C×35s; annealing: 60°C × 35s; and elongation: 60°C×40s, for 40 cycles. PCR reactions were performed in triplicate using 20 µL reaction mixture containing 4 µL of 5×FastPfu buffer, 2 µL of 2.5 mM dNTPs, 0.8 µL of each primer (5µM), 0.4 µL of FastPfupolymerase, and 10ng of template DNA. According to the formula, the relative mRNA expression level of each sample has been expressed as 2^−ΔΔCT^, Ct= target gene Ct value-β-actin Ct value, and the data obtained were analyzed. The primer sequences of specific genes investigated are as follows:

**Table 1.**
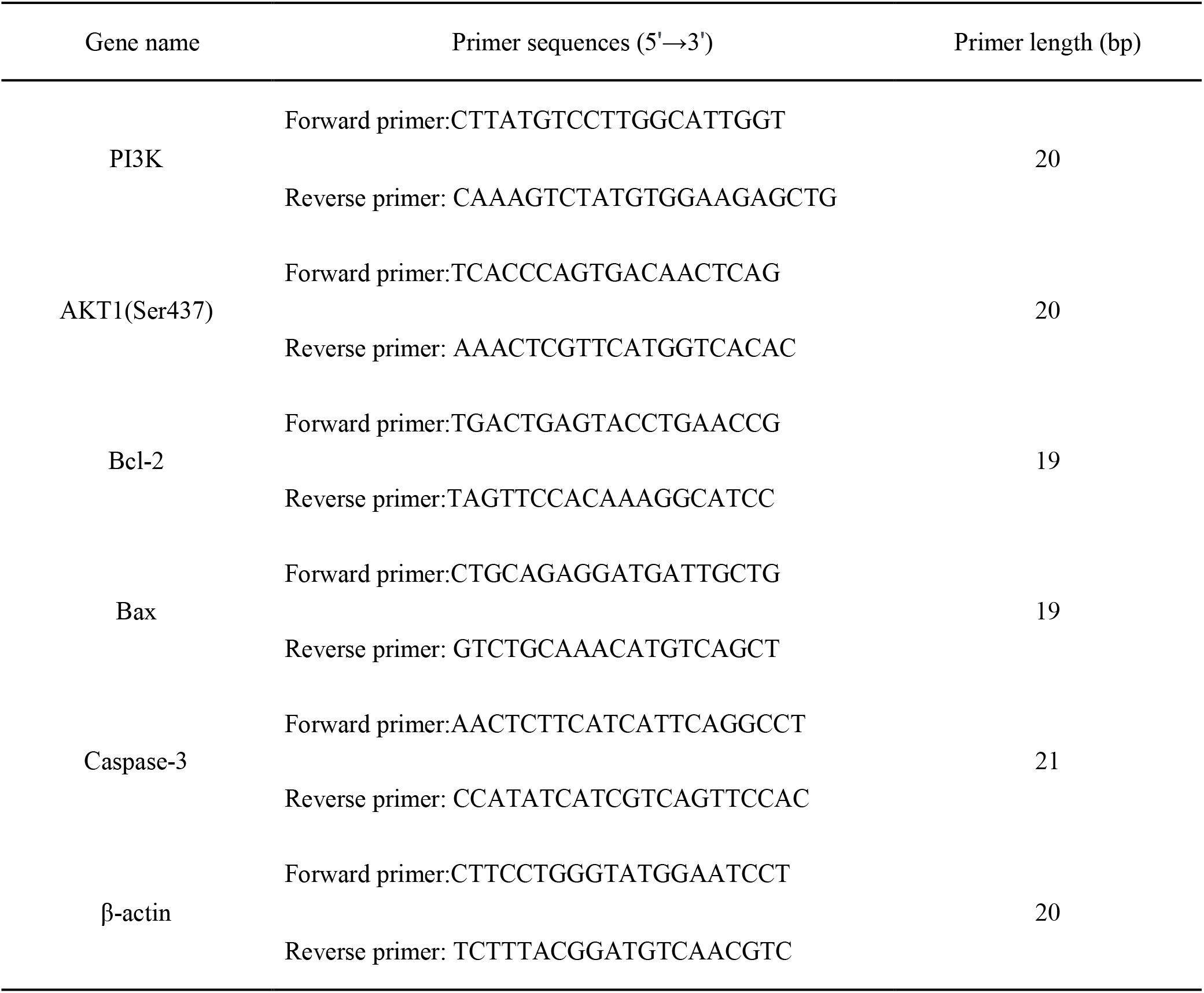
Primer sequences.

### 4.15 Immunofluorescence labeling of VSMCs

VSMCs were subjected to fixation with 4% paraformaldehyde in 0.1 MPBS for 30 min. Following subjection to rinsing steps with PBS, the coverslips with adherent cells were used for immunofluorescence staining. The steps for immunofluorescence staining of paraffin-embedded sections were as follows: the sections were dewaxed in xylene to water, followed by immersion of the sections in sodium citrate buffer; sections were heated in a microwave oven for approximately 20 min to perform antigen retrieval, followed by subjection to washing steps thrice in PBS, with each stepspanning 5 min. After the sections were almost dried, 10% BSA was added and the section were incubated at room temperature for 35 min. The blocking solution was gently washed off, and a primary antibody was added that had been formulated in a certain proportion to the sections. Bax, Bcl-2 (1:200), caspase-3, mTOR (1:100), α-SMA, SM22α, and OPN(1:250)were added to the sections in a wet box, following which incubation was performed at 4°C overnight. After subjection to rewarming for 35 min, the sections were washed 3 times in PBS, with each step spanning 5 min. When the sections were slightly dried, the corresponding fluorescent secondary antibody was added to cover the tissues completely and incubation was performed in the dark for 60 min. After washing the sections 3 times in PBS, 200 μL DAPI was added to the sections and incubation was conducted at room temperature for 10 min. The sections were washed 3 times in PBS again. Subsequently, the sections were dried, and an appropriate amount of an anti-fluorescent quencher resin mounting tablet was added in a dropwise manner and results were observed under a microscope.

### 4.16 Statistical analysis

The data have been presented as mean ± SD values. We performed the LSD test in the one-way ANOVA module of the SPSS 25.0 software (IBM, SPASS Statistics 25.00) to perform pairwise comparisons between multiple groups. Statistical graphs were generated using the GraphPad Prism 7.0 software (GraphPad, Inc., La Jolla, CA, USA). A P value <0.05 was considered to indicate statistical significance. The duplicate or triplicate values derived from as ingle experiment were averaged.

## Study approval

All processes involving animals were authorized by the Animal Research Ethics Committee of Kunming Medical University (Animal Ethics License Number KMMU2018219).

## Supporting information

Cover letter

Manuscript

## Authors’ contributions

Author contributions: L. Zhang, Z. Yu, H.T. Qin, and P. Chen designed research; L. Zhang, Y.T. Tao, Q. Hu, R.H. Yang, J. Jia, M.Y. Yu, D. YU, Y.T. Wang, J.N. Shi, H.N. Jin, Y. Yang, Y.Z. Yang, G.P. Tang, J. Xu, B. Xiong, Z.Q. Shen, Z. Yu, H.T. Qin, and P. Chen performed research; L. Zhang, Y.T. Tao, Q. Hu, R.H. Yang, J. Jia, M.Y. Yu, Z. Yu, H.T. Qin, and P. Chen analysed data; L. Zhang, Y.T. Tao, Q. Hu, Z. Yu, H.T. Qin, and P. Chen wrote the paper; L. Zhang and P. Chen secured funding.

## Acknowledgments

Acknowledgement: This work was supported by the National Natural Science Foundation of China [grant numbers 81860641 and 81260493], the Key Project of Natural Science Foundation of Yunnan Province, P.R. China [grant number 202001AS070035], and the Innovation Foundation for Doctoral Students of Kunming Medical University [grant number 2021D05].

