## Supplementary material for "A novel mechanism of Euonymine inhibits in-stent restenosis through enhancing contractile phenotype of VSMCs by targeting AKT1 and p38MAPK": Cover letter

October 29, 2021

Dear Editor:

We wish to submit a research paper for publication in the *The journal of Clinical Investigation* titled “ Euonymine inhibits ISR through enhancing contractile phenotype of VSMCs via Targeting AKT1 and p38MAPK”.

*Tripterygium hypoglaucum (Levl) Hutch* (THH) is a traditional Chinese medicine that has been used therapeutically for rheumatoid arthritis, nephrotic syndrome, and systemic lupus erythematosus. Euonymine(C_38_H_47_NO_18_, 805.783), a crucial monomer active constituent extracted from THH, have drawn wide attention of scholars at home and abroad because of its important medicinal value, unique pharmacological effects. In recent years, we have studied on cardiovascular activity of Euonymine and we found that Euonymine inhibited VSMCs phenotypic transformation by targeting AKT1 to regulate PTEN/AKT1/mTOR signaling pathway, exerting anti-proliferative, anti-migratory and pro-apoptotic effects on ox-LDL-induced cell injury model. In addition, we demonstrated that Euonymine induced apoptosis of VSMCs via the p38MAPK-related mitochondria-dependent apoptotic pathway. Herein, we applied the rabbit carotid balloon injury and porcine atherosclerotic coronary implantation models to simulate the dynamic pathological progression of ISR after PCI in humans. The dynamic development process of ISR in these two animal models is close to the clinical pathology of ISR after PCI, which has valuable clinical reference value. In addition, we evaluated the effect of Euonymine on post-PCI ISR by QCA and OCT techniques in a preclinical porcine coronary artery stent model. The results corroborated each other to visualize the extent of lesions in vascular neointimal hyperplasia, and visually confirmed that coronary intimal hyperplasia was effectively suppressed on the ISR stent model after implantation of Euonymine coated stents, and showed good histocompatibility and safety. Therefore, Euonymin is a potential stent-coated for treating ISR after PCI.

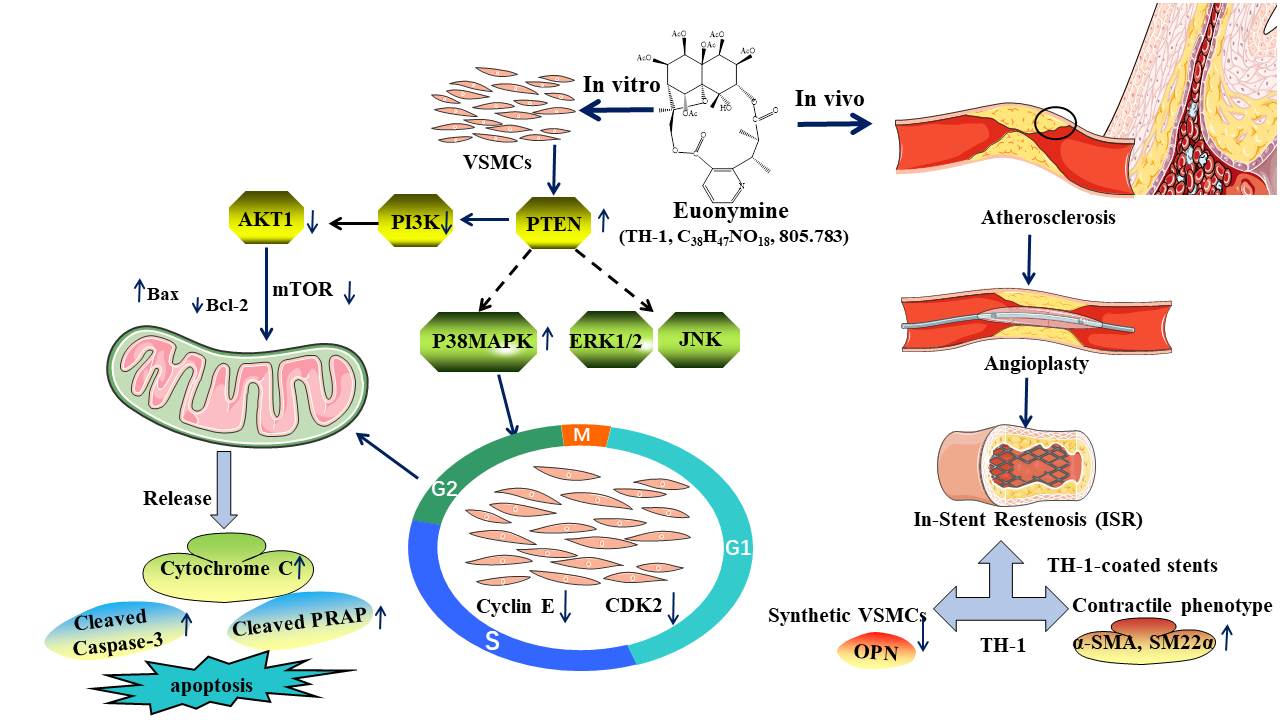

Figure.1 Graphical Abstract

Further, we believe that our research will be of interest to the readership of *the journal of Clinical Investigation*, and we provide mechanistic insight into the effects and mechanism of a plant-based monomer on the prevention and treatment of ISR after PCI.

This manuscript has not been published or presented elsewhere in part or in entirety and is not under consideration by another journal. The study design was approved by the appropriate ethics review board. We have read and understood your journal’s policies, and we believe that neither the manuscript nor the study violates any of these. There are no conflicts of interest to declare.

Thank you for your consideration. I look forward to hearing from you.

Sincerely yours,

PengChen, Ph.D.

Professor.

School of Pharmaceutical Sciences and Yunnan Key Laboratory of Pharmacology for Natural Products, Kunming Medical University, Kunming, 650500, China.
