## Supplementary material for "A novel mechanism of Euonymine inhibits in-stent restenosis through enhancing contractile phenotype of VSMCs by targeting AKT1 and p38MAPK": Manuscript

**4.1 Chemicals and reagents**

α-SMA and OPN were purchased from Boster Biological Technology, Ltd. (Wuhan, China), and anti-SM22α antibodies were purchased from Santa Cruz Biotechnology, Inc. (Santa Cruz, CA, USA). Anti-Caspase-3 antibody, anti-PI3K antibody, anti-p-PI3K antibody were purchased from Abcam Company(Shanghai, China); anti-AKT antibody, anti-Phospho-AKT1(Ser473)antibody, anti-mTOR antibody, and anti-Phospho-mTOR(Ser2448)antibody were purchased from American CST Company; anti-Bcl-2 Antibody was purchased from Boster Biological Technology Co.Ltd.; anti-Bax antibody was purchased from Affinity Biosciences. The BCA protein quantification kit was purchased from BiyunTian Company (Shanghai,China), the ECL luminescence reagent was purchased from Millipore(Shanghai, China), the total RNA extraction kit was purchased from TIANGEN(Beijing, China), and the total RNA extraction kit was purchased from TIANGEN. The Revert Aid First SndcDNA Synthesis Kit was purchased from ThermoFisher(Shanghai, China), and the RT-PCR MasterMix was purchased from Vazyme (Nanjing, China). 5% pentobarbital sodium was purchased from the First Hospital of Beijing University, and monoclonal anti-PCNA antibody, universal SP immunohistochemical detection kit, and DAB were purchased from Beijing Zhongshan Biotechnology Co. Ltd. MK-2206, SB203580, and SP600125 were purchased from Selleck.cn; the required solution concentration was prepared according to the manufacturer's instructions. Bayaspirin(aspirin enteric-coated tablets) was purchased from Bayer Medical Care Co. Ltd. (Germany); Plavix (clopidogrel) was purchased from Sanofi-aventis Pharmaceuticals LTD (France); pentobarbital injection was purchased from Shanghai Jingrong Biological Technology Co. Ltd. (China); and serazine hydrochloride injection was purchased from Jilin Founder Animal Pharmaceutical Co. Ltd. (China).

**4.3.3** **Establishment of porcine coronary artery stent model, sample collection, and pathological examination**

**4.3.3.1 Establishment of porcine coronary artery stent model**

I. Animal grouping: Twenty minipigs were randomly divided into Bare metal stent (BMS), three Euonymine-eluting Stent groups (0.14 [low-dose], 0.70 [medium-dose], and 2.80 [high-dose] μg/mm^2^), and Sirolimus-eluting Stent (0.14μg/mm^2^) group. A model of porcine atherosclerosis was replicated according to the previous research of our research team[25], and then a stent was placed in the coronary stenosis site, and left or right coronary angiography and OCT were performed through the right femoral artery or left femoral artery route after stent implantation and 28 days，(See later for details of the specific operation).

II. Pre-operative preparation：All surgical animals were prepared before surgery and fasted with no food and water on the morning of surgery. Before surgery, pentobarbital sodium salt injection (0.1 ml/kg) was administered intravenously, while xylazine hydrochloride injection (0.01 mL/kg) was administered intramuscularly for general anesthesia. Additional anesthetics were administered as necessary during surgery. The experimental animals were then prepared for preoperative skin preparation and intraoperative continuous cardiac monitoring.

III. Femoral artery puncture or femoral artery dissection: The porcine femoral artery was punctured with a 12-gauge needle 1~1.5 cm below the inguinal ligament using the Seldinger method and a 6F arterial sheath was placed. After successful puncture, a guide wire was inserted, and the 6F arterial sheath was inserted into the femoral artery along the guide wire, and the arterial sheath was sutured and fixed.

IV. Coronary angiography and stent implantation: Heparin 200 lU/Kg was administered through the sheath for systemic heparinization, ensuring ACT >300 seconds. The 6F Juking Left 3.5 guiding catheter was delivered to the aortic sinus via the contrast guidewire under the left anterior oblique 30° trans-x-ray fluoroscopy, the contrast guidewire was withdrawn, the triple-ring syringe and pressure set were attached, the guiding catheter was adjusted to the left coronary artery orifice, the left coronary angiogram was performed, and the 0.014-inch Runthrough or BMW guidewire was delivered to the distal end of the anterior descending branch or gyrus branch, and the flat and straight segment with few branches and suitable vessel diameter was selected as the stent placement site. The stent is then released at a ratio of 1:1.1 to 1.2 between the vessel and the stent, and the stent release pressure is usually 10 atm ×12 seconds. The stent balloon was measured and coronary angiography was performed again to confirm lumen patency, antegrade flow to TIMI level 3, no filling defect, wall entrapment, thromboembolism, stent displacement, etc. Qualitative OCT was performed to further evaluate the stent apposition, endothelial tear and tissue prolapse. (See later for details of the OCT procedure). Right coronary artery stent placement and OCT catheterization were performed in the same way.

V. Surgical puncture site handling: After the surgery is completed, the OCT catheter, guidewire and guiding catheter are withdrawn successively, and the arterial sheath is removed. In the case of inguinal skin incision to expose the femoral artery for puncture, the skin is sutured layer by layer and then the arterial sheath is removed, and pressure is applied to the puncture site for 25-30 minutes until there is no bleeding and oozing at the puncture site.

VI. Postoperative treatment: The resuscitation of the animal was personally monitored by the researcher for the first 1-2 hours after the catheterization operation, and the vital signs were monitored during the anesthesia awakening period of the experimental animal. Postoperatively, penicillin 4 million units was given intramuscularly once daily for three days, heparin sodium 6000 IU was given subcutaneously twice daily for 1 week, and clopidogrel (75 mg/d) and aspirin (100 mg/d) were administered orally for 28 days.

VII. Coronary angiography and analysis

The preoperative preparation was basically the same as that of the previous coronary stenting procedure. A 6F arterial sheath was delivered via femoral artery puncture, and a 6F guiding catheter was sent to the coronary artery port for coronary angiography(FD10/10，Philip Co.Ltd)，Germany) via a contrast guidewire, and the presence of significant stenosis or occlusion was assessed by multiple body angiography crude rates. Quantitative coronary angiography (QCA) is performed as close to the proximal end of the lesion as possible, avoiding major bifurcations and openings, and within 10mm of the measurement point. The relative vessel internal diameter, the minimum internal diameter in the stent, and the degree of stenosis were measured by a combination of QCA software analysis and visual inspection.

VIII. OCT examination and analysis

Deliver the Runthrough guidewire to the distal coronary artery of the stent placed in the same way as for stent placement. Flush the manifold with 5 ml of pure iodine contrast and remove all air from the manifold until 3-5 drops of contrast flow from the end tip of the catheter, align the white connector with the DOC connector and rotate clockwise 1/8 of a turn to lock it in place. Once the OCT imaging system is ready, the Dragonfly imaging catheter is sent along the guidewire to the guiding catheter for recalibration, and under fluoroscopic guidance, the catheter is moved forward until the proximal marker (lens After the Dragonfly imaging catheter was injected into the lumen of the vessel with a syringe pump, the lumen was flushed to remove blood from the target vessel, and the imaging guide wire was automatically retracted at a speed of 1.0 mm/s. The imaging was completed with a video monitor at 15.6 frames/s. OCT imaging and analysis are performed with the C7XR OCT imaging system (St. Jude Medical Co.Ltd).

OCT imaging analysis: quantitative analysis was performed by applying the software within the OCT system after the imaging was completed, and the most narrowed area within each stent was selected to measure the neointimal thickness (mm): the distance between the neointima on the surface of the lumen and the surface of the stent column was the neointimal hyperplasia thickness, measured as perpendicular to both as possible; residual lumen area (mm^2^): the actual area of the vascular lumen: the outer boundary was the intima; stent area ( mm^2^): the total area contained was measured with the stent as the boundary; area of neointima (mm^2^) = the total area between the inner elastic membrane as the outside and the inner membrane as the inner boundary, i.e. the area of hyperplasia within the stent; degree of stenosis = area of neointima/cross-sectional area of the inner elastic membrane.
